# Transcriptome analysis for the development of cell-type specific labeling to study olfactory circuits

**DOI:** 10.1101/2020.11.30.403865

**Authors:** Anzhelika Koldaeva, Cary Zhang, Yu-Pei Huang, Janine Reinert, Seiya Mizuno, Fumihiro Sugiyama, Satoru Takahashi, Taha Soliman, Hiroaki Matsunami, Izumi Fukunaga

## Abstract

In each sensory system of the brain, mechanisms exist to extract distinct features from stimuli to generate a variety of behavioural repertoires. These often correspond to different cell types at some stage in sensory processing. In the mammalian olfactory system, complex information processing starts in the olfactory bulb, whose output is conveyed by mitral and tufted cells (MCs and TCs). Despite many differences between them, and despite the crucial position they occupy in the information hierarchy, little is known how these two types of projection neurons differ at the mRNA level. Here, we sought to identify genes that are differentially expressed between MCs and TCs, with an ultimate goal to generate a cell-type specific Cre-driver line, starting from a transcriptome analysis using a large and publicly available single-cell RNA-seq dataset (Zeisel et al., 2018). Despite many genes showing differential expressions, we identified only a few that were abundantly and consistently expressed only in MCs. After further validating these putative markers using *in-situ* hybridization, two genes, namely *Pkib* and *Lbdh2*, remained as promising candidates. Using CRISPR/Cas9-mediated gene editing, we generated Cre-driver lines and analysed the resulting recombination patterns. This analysis indicated that our new inducible Cre-driver line, *Lbhd2-CreERT2*, can be used to genetically label MCs in a tamoxifen dose-dependent manner, as assessed by soma locations, projection patterns and sensory-evoked responses. Hence this line is a promising tool for future investigations of cell-type specific contributions to olfactory processing and demonstrates the power of publicly accessible data in accelerating science.

## Introduction

The complexity of the brain, in part, originates from the diversity of its components, the rich variety of cells. This diversity is evident in morphology, connectivity, molecular expression profiles, and biophysical properties (Luo et al., 2018; Sanes and Masland, 2015; Zeng and Sanes, 2017), which together give rise to what we refer to as cell types. Because the differences are thought to reflect distinct computational tasks or functions (Luo *et al.*, 2018; Masland, 2004), the ability to selectively identify, and to manipulate, each cell type experimentally is key to understanding how the brain works.

In rodents, complex, synaptic processing of olfactory information in the brain first occurs in the olfactory bulb. The principal cells of the olfactory bulb, the MCs and TCs, convey the output of this region and are thought to form parallel information streams. They differ in a variety of anatomical and physiological properties (Economo et al., 2016; Fukunaga et al., 2012; Igarashi et al., 2012; Jordan et al., 2018; Kapoor et al., 2016; Otazu et al., 2015; Phillips et al., 2012). MCs, which are the larger of the two, are thought to form distinct circuits with local neurons from those formed by TCs (Fukunaga *et al.*, 2012; Geramita et al., 2016; Mori et al., 1983; Phillips *et al.*, 2012), some of which may explain the differences in how they encode odors. For example, in TCs, the timing of responses adheres strictly to a specific phase of the sniff cycle, while MCs modulate the timing widely over the entire sniff cycle (Ackels et al., 2020; Fukunaga *et al.*, 2012; Igarashi *et al.*, 2012). Signal integration over this long temporal window is thought to allow MCs to represent more complex information (Fukunaga *et al.*, 2012). Further, in contrast to TCs whose axons project to a more limited portion of the olfactory cortex, the target areas of MCs range widely, extending as far as the posterior piriform cortex, the cortical amygdala, and the lateral entorhinal cortex (Haberly and Price, 1977; Igarashi *et al.*, 2012), indicative of a variety of behavioural contexts in which MCs are likely to be important.

Despite the fundamental roles these two cell types play in olfaction, suitable molecular markers and genetic tools are lacking. Molecules commonly used to label the output neurons of the olfactory bulb include protocadherin-21 (Nagai et al., 2005), T-box transcription factor 21 (*Tbx21*, also known as *Tbet*; (Faedo et al., 2002; Papaioannou and Silver, 1998)), as well as cholecystokinin (*Cck*) (Seroogy et al., 1985). In the brain, Tbx21 is expressed from embryonic day 14 (Faedo *et al.*, 2002) and is exclusive to the principal neurons of the olfactory bulb, labeling both MCs and TCs (Faedo *et al.*, 2002; Haddad et al., 2013; Mitsui et al., 2011). In contrast, *Cck* is expressed widely in the brain (Larsson and Rehfeld, 1979; Taniguchi et al., 2011). In the olfactory bulb, expression occurs preferentially in TCs over MCs (Seroogy *et al.*, 1985), which has been utilized for analysing the unique physiology of TCs (Economo *et al.*, 2016; Short and Wachowiak, 2019). These overlapping but differential expression patterns between *Cck* and *Tbx21* may be useful in discovering more selective markers to distinguish the two types of principal neurons.

A variety of methods now exist to analyse gene expression patterns in relation to cell types, including *in-situ* hybridization (Lein et al., 2006), as well as transcriptomic approaches that retain spatial information, with increasing resolution (Ståhl et al., 2016). More recently, single-cell RNA sequencing (scRNA-seq) (Pfeffer et al., 2013; Shekhar et al., 2016; Sugino et al., 2006; Tang et al., 2009; Tasic et al., 2016; Zeisel et al., 2015) has seen rapid developments, which have enabled the investigation of cell-type specific gene expression patterns with unprecedented level of detail and scale (Lein et al., 2017; Zeng and Sanes, 2017). A useful application of this information in turn may be to generate transgenic driver lines, that allow a particular cell type to be extensively studied. The availability of Cre-driver lines has been instrumental in revealing unique functions of distinct cell types, across multiple levels of analyses (Cruz-Martín et al., 2014; Daigle et al., 2018; Dhande et al., 2013; Gong et al., 2003; Madisen et al., 2015; Madisen et al., 2012; Sanes and Masland, 2015; Taniguchi *et al.*, 2011; Wolff et al., 2014).

Here, we take advantage of a large dataset that has become publicly available (Zeisel et al., 2018), to discover markers that distinguish between MCs from TCs. The results of the analyses allowed us to generate, and characterize, new Cre-driver lines. Such molecular tools will be key to understanding the mechanisms of olfactory perception and behaviour.

## Results

In search of molecular markers, we sought to compare the gene expression patterns of MCs and TCs. This may reveal candidate markers, which are genes that are selectively enriched in the target cell-type of interest, in this case MCs, but not TCs. This first requires a method to identify MCs and TCs in a gene expression data, and, second, distinguish their gene expression profiles from each other. Previous studies observed that *Tbx21*, a T-box type transcription factor, labels both MCs and TCs (Faedo *et al.*, 2002; Haddad *et al.*, 2013; Mitsui *et al.*, 2011), while the neurotransmitter cholecystokinine (*Cck*) is more abundant in TCs (Economo *et al.*, 2016; Seroogy *et al.*, 1985). To verify these distributions in our hand, we crossed *Tbet-Cre* and *Cck-IRES-Cre* lines (Haddad *et al.*, 2013; Taniguchi *et al.*, 2011) with the Rosa-CAG-LSL-tdTomato reporter line, *Ai14* (Madisen *et al.*, 2012), for Cre-dependent expression of the red fluorescent protein, tdTomato. We confirm that Tbx21-driven expression labels cells in the mitral cell layer and the external plexiform layer where TCs are located, while *Cck*-driven expression labels a larger number of cells all over the olfactory bulb (Fig. 1A,B, Supplementary Fig. 1), especially those that extend more superficially in the glomerular layer and sporadically in the granule cell layer. Importantly, labeling coupled to *Cck* expression is less consistent in cells that occupy the mitral cell layer. This differential expression patterns between *Tbx21* and *Cck* may be used to distinguish MCs from TCs in gene expression data (Fig. 1B,C).

**Figure 1:**
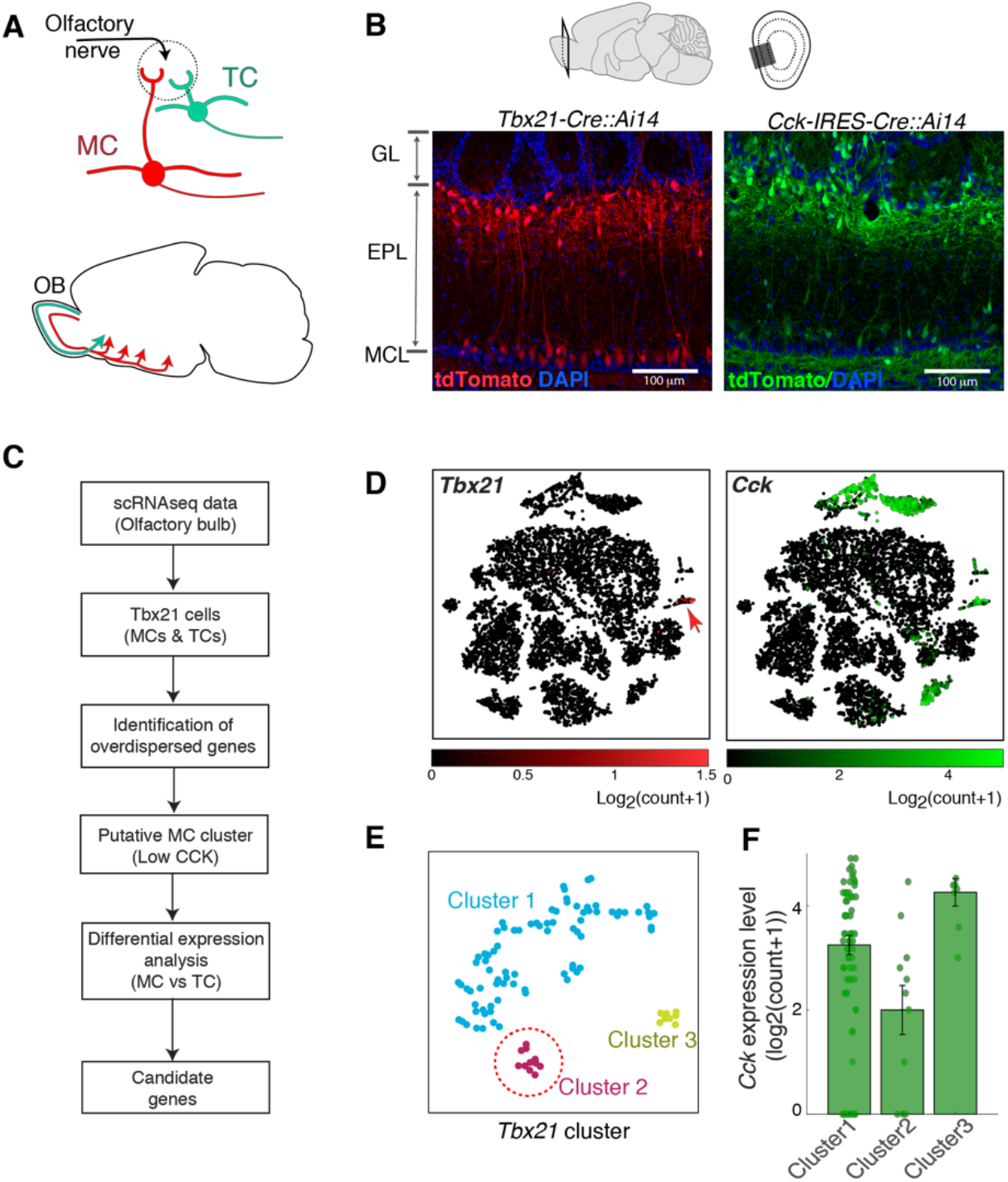
Strategy for identifying mitral-specific markers from scRNA-seq data. (**A**) Schematic showing major anatomical differences between the two cell types; MCs (red) are located deeper in the olfactory bulb (OB) layers, and project widely in olfactory cortices. TCs (green) are smaller, superficially located principal neurons that project to anterior portions of the olfactory cortex. (**B**) Tbx21 and CCK expression patterns in the main olfactory bulb; example images showing tdTomato expression patterns in Tbet-cre∷Ai14 mouse (red) and Cck-IRES-cre∷Ai14 mouse (green). Scale bar = 100 μm. GL = glomerular layer, EPL = external plexiform layer, MCL = MC layer. (C) Schematic of workflow; Putative mitral cluster from scRNA-seq data is identified by the observation that MCs and TCs both express *Tbx21*, but *Cck* is more abundant among TCs. Once a MC and TC clusters are identified, differential expression analysis is carried out to identify genes that are selectively expressed in MCs. (**D**) OB cells plotted in tSNE coordinates, with *Tbx21* and *Cck* expression levels (left and right panels, respectively) indicated with colour maps shown below. (**E**) *Tbx21*-positive cluster was further analysed and the sub-clustered and displayed in new tSNE coordinates. (**F**) *Cck* expression levels for the sub-clusters in (**E**). Cluster 2 has the lowest level and is inferred as the putative MC cluster (red dotted line in **E**).

Identification of molecular markers by differential expression analyses requires a robust and large dataset, especially when distinguishing similar cell types, such as in the case of MCs and TCs. We turned to a public, large scale scRNA-seq dataset of the mouse brain (Zeisel *et al.*, 2018). This contains data from approximately 0.5 million cells, 10,745 cells of which are from the olfactory bulb. We clustered the data based on the similarity of gene expression patterns. To achieve this efficiently, we identified top 500 overdispersed genes out of 27,998 genes in the dataset (see Methods). Such genes are highly informative for determining genetic differences among the cells. Using this reduced dataset, we performed Principal Component Analysis (PCA), followed by t-distributed Stochastic Neighbor Embedding (tSNE) on the first 10 principal components to further reduce the dimensionality of the gene expression space to two. The combination of the two algorithms preserves both the global and local structures of the data (Kobak and Berens, 2019). To obtain clusters, hierarchical density-based special clustering algorithm (HDBSCAN (Campello et al., 2013)) was applied on the 2-dimentional tSNE space to cluster the data (see Methods). Within the olfactory bulb dataset, we found that 1,682 cells belong to *Cck*-positive clusters, while *Tbx21-*expressing cluster comprised 101 cells. Generally, expression patterns of *CCK* and *Tbx21* together mirror those of *Slc17a7* (VGlut1) and Slc17a6 (Vglut2), indicating that they are mainly glutamatergic populations (Supplementary Fig.2), with the largest portion of glutamatergic, *Cck*-positive clusters outside of the Tbx21-positive cluster. Further, a small set of *Cck*-expressing neurons did not overlap with *Slc17a7*- & *Slc17a6*-positive clusters (Supplementary Fig. 2). To identify a putative mitral cluster from scRNA-seq data, we took advantage of the observation that MCs and TCs both express *Tbx21*, but *Cck* is more abundant among TCs. In the *Tbx21*-positive cluster, the second largest cluster (cluster 2, Fig. 1E) showed the lowest *Cck* expression level (Fig. 1F). We thus refer to this as the putative MC cluster, and refer to the remaining as TC cluster (TC1).

An ideal molecular marker should be expressed abundantly and consistently in the cell type of interest, while having minimal expression levels in other cell types. To search for candidates with these properties, gene expression patterns of putative MCs were compared against the rest of *Tbx21*-expressing neurons (TC1; Fig. 2A), as well as glutamatergic, *Cck*-positive clusters outside of the *Tbx21*-cluster (TC2; Fig. 2A). First, Mann-Whitney U-test was used to screen genes that are differentially expressed, with p-values adjusted using the Benjamini-Hochberg procedure. This procedure identified several differentially expressed genes (Table 1), at the adjusted p = 0.05 level. Among these were *Calb2* (calbindin 2 gene), *Ntng1* (netrin G1), *Ppm1j* (protein phosphatase 1J), *Rph3a* (rabphilin 3A), *Kcnq3* (voltage-gated potassium channel subfamily Q member 3), and *Chrna2* (nicotinic cholinergic receptor alpha). Of the differentially expressed genes, we focused on those that are present in the majority (>50%) of cells in the putative MC cluster, but in less than 10% of the cells outside of this cluster (Fig. 2B; Table 1). Only a small number of the differentially expressed genes fulfilled these criteria, and even fewer showed minimal expression levels outside of MCs, as judged by the OB-wide expression patterns (Supplementary Fig 3), as well as by the *in situ* hybridization data in the Allen Brain Atlas (Lein et al., 2007). Candidate genes that showed clear hybridization signals outside of the mitral cell layer were therefore not pursued further (Supplementary Fig. 3).

**Figure 2:**
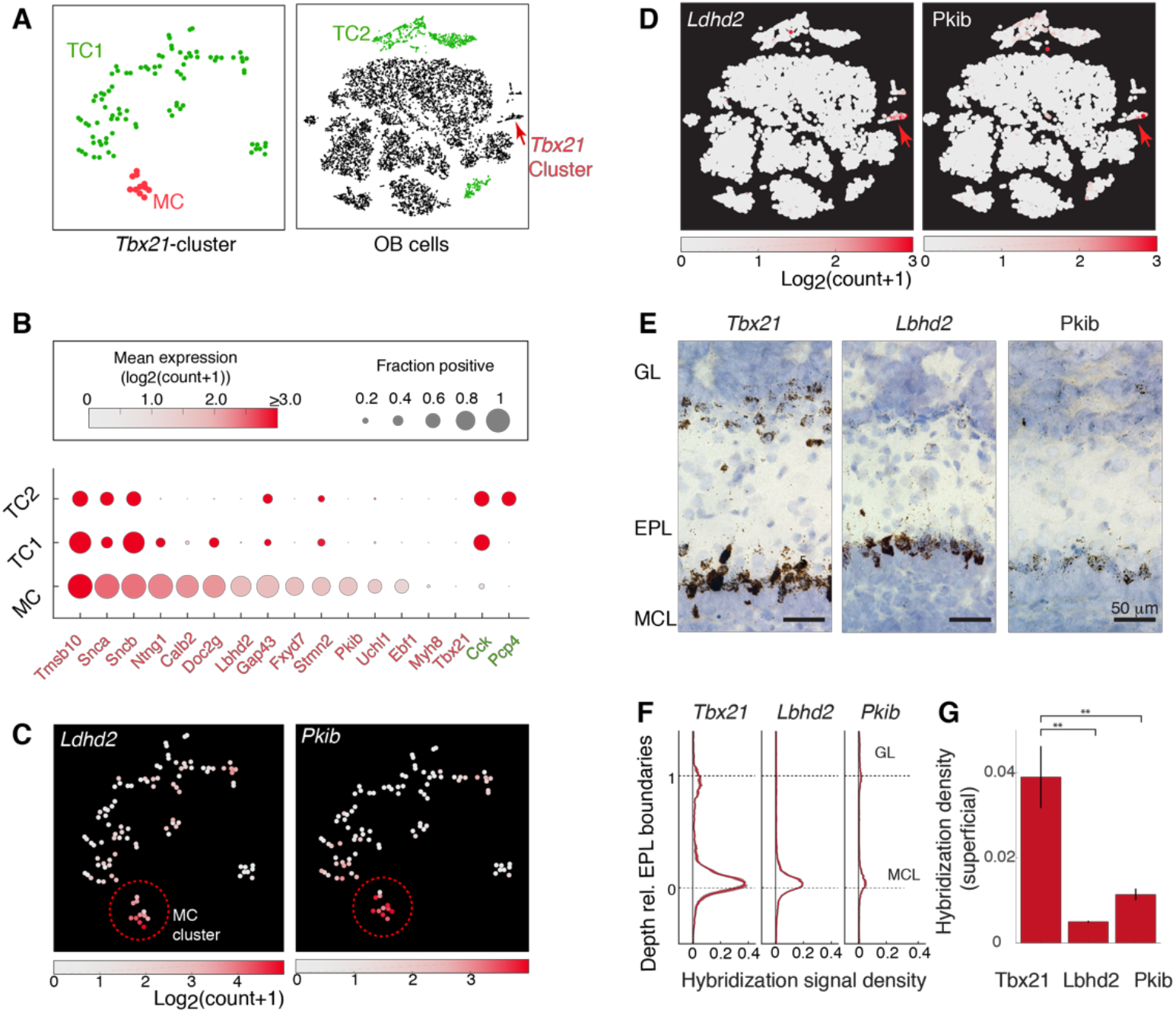
Differential gene expression analysis reveals candidate marker genes for MCs. (**A**) Cluster grouping used for differential gene expression analysis to distinguish TCs from MCs. *Tbx21* expressing cells (left) constitute the MC cluster (red) and TC1 cluster (green), which is identical to the combined clusters 1 and 3 shown in Fig. 1E. The olfactory bulb (OB)-wide dataset (right) contains the TC2 cluster (green) which is equivalent to the *Cck*-rich clusters shown in Fig. 1D without the *Tbx21*-rich clusters. (**B**) Genes that are significantly enriched in MCs (red), as well as those that are enriched in TC clusters (green). The size of data points indicates the consistency of expression, measured as the fraction of cells in the cluster that express the gene. Mean expression level (log2(count+1)) is color-coded as shown in the colormap above. (**C**) Expression levels of two candidate genes, *Pkib* and *Lbhd2*, with the corresponding colormaps, superimposed on the 3-sub clusters of the *Tbx21*-rich cluster (same tSNE coordinate as Figure 1E) and (**D**) the whole OB data. Red arrow points to the *Tbx21*-cluster. (**E**) Example *in-situ* hybridization signals revealed by DAB staining, for *Tbx21* (left), *Lbhd2* (middle) and *Pkib* (right) for the MOB layers indicated. Scale bar = 50 μm. (**F**) *In-situ* hybridization signal density relative to the external plexiform layer boundary (0 – 1); hybridization signal was thresholded, and the proportion of pixels above the threshold for each normalized external plexiform layer depth was expressed as density. (**G**) Summary of hybridization signals in the superficial locations (depth upper half of external plexiform layer). N = 3 mice, with samples from dorsal, ventral, medial and lateral locations at middle and caudal levels of the antero-caudal axis.

**Table 1:** List of genes differentially expressed between mitral cells and tufted cells.

| Gene Name | Mean Expression<br>(log2(count+1)) | Adjusted p value |
| --- | --- | --- |
| Myh8 | 2.14 | 1.57E-09 |
| Pkib | 3.40 | 7.02E-06 |
| Fxyd7 | 3.66 | 1.41E-05 |
| A230065H16Rik (Ldbh2) | 3.77 | 1.55E-05 |
| Ebf1 | 2.55 | 1.55E-05 |
| Calb2 | 4.60 | 1.55E-05 |
| Snca | 5.10 | 1.55E-05 |
| C1ql1 | 2.53 | 1.55E-05 |
| Cxcl14 | 0.42 | 1.55E-05 |
| Sostdc1 | 0.36 | 2.07E-05 |
| Tmsb10 | 6.12 | 3.05E-05 |
| Spp1 | 2.20 | 3.45E-05 |
| Ntng1 | 4.76 | 4.25E-05 |
| RP23-407N2.2 | 1.81 | 4.70E-05 |
| 1110008P14Rik | 2.45 | 0.000163822 |
| Tspan17 | 1.99 | 0.000211554 |
| Uchl1 | 3.24 | 0.000253744 |
| Shisa3 | 1.98 | 0.000253744 |
| Gap43 | 3.69 | 0.000439148 |
| Ppm1j | 1.85 | 0.000449468 |
| Rph3a | 0.74 | 0.000449468 |
| Crtac1 | 1.97 | 0.000449468 |
| Tnnc1 | 0.27 | 0.000457871 |
| Mmp17 | 0.45 | 0.000585813 |
| Nov | 0.18 | 0.000707168 |
| Gng13 | 1.03 | 0.001406527 |
| Rab15 | 1.62 | 0.001959665 |
| Gm27199 | 1.11 | 0.001978521 |
| Vsnl1 | 0.83 | 0.002521565 |
| Stmn2 | 3.57 | 0.003524909 |
| Npr1 | 1.30 | 0.003848843 |
| Doc2g | 4.52 | 0.004924184 |
| Nptxr | 0.78 | 0.005760704 |
| Kcnq3 | 0.94 | 0.005928465 |
| Slc1a2 | 1.29 | 0.007685702 |
| Sncb | 4.97 | 0.00781742 |
| Ephx4 | 1.37 | 0.009259688 |
| Atp9a | 1.91 | 0.009383075 |
| Nrsn1 | 2.65 | 0.01005383 |
| Diras2 | 1.50 | 0.01094975 |
| Rgs4 | 0.76 | 0.01098611 |
| Al413582 | 2.09 | 0.0124281 |
| Abi3bp | 0.09 | 0.01393499 |
| Lingo1 | 1.67 | 0.01393499 |
| Tshz2 | 3.22 | 0.01393499 |
| Mal2 | 0.18 | 0.01393499 |
| Fkbp1b | 0.00 | 0.01393499 |
| Prkcb | 1.23 | 0.01504482 |
| Cdh4 | 1.02 | 0.01504482 |
| Meg3 | 7.01 | 0.01660475 |
| Chrna3 | 0.61 | 0.01660475 |
| Ifitm10 | 0.74 | 0.01905793 |
| Adk | 0.34 | 0.02020242 |
| Stx1a | 0.89 | 0.02020242 |
| Igfbp5 | 2.75 | 0.02381658 |
| Fam19a1 | 0.09 | 0.0261028 |
| Grin1os | 0.09 | 0.0261028 |
| Cdc20 | 0.09 | 0.0261028 |
| Gm26803 | 0.09 | 0.0261028 |
| Resp18 | 1.14 | 0.0261028 |
| Pantr1 | 1.12 | 0.02678842 |
| Lmo4 | 0.91 | 0.02690157 |
| Pvrl1 | 1.07 | 0.03585761 |
| Kitl | 0.00 | 0.0397583 |

Based on the initial screening, *Pkib* (protein kinase inhibitor beta) and *Lbhd2* (LBH domain containing 2; Fig. 2C,D) genes fit the criteria for the MC marker (Fig. 2C). To confirm that these genes indeed are selectively expressed in MCs, we carried out *in-situ* hybridization for *Pkib* and *Lbhd2* (Fig. 2B–D) on olfactory bulb sections. Indeed, probes for *Pkib* and *Lbhd2* gave rise to monolayer-like signals at the lower boundary of the external plexiform layer, corresponding to the location of the mitral cell layer. For quantification, *Pkib* and *Lbhd2* signals were expressed as density (Methods) and plotted relative to the boundaries of the external plexiform layer. This revealed that *Pkib* and *Lbhd2* both label cells in the MC layer, with significant reduction in the superficial signals corresponding to TCs, especially compared to *Tbx21* (Fig. 2D; mean signal densities in the upper external plexiform layer: *Tbx21* = 0.39 ± 0.007; *Lbhd2*= 0.005± 0.0002; *Pkib* =0.01± 001, p <0.001 1-way ANOVA, F = 17.9, degrees of freedom = 2). Hybridization signal in the MCL was relatively uniform throughout the olfactory bulb (Supplementary Fig. 4), while residual expression patterns of *Pkib* and *Lbhd2* in non-MC cells differed somewhat, with faint signals in the glomerular layer and external plexiform layer for *Pkib* and *Lbhd2*, respectively. Thus, *Pkib* and *Lbhd2* are promising candidates for selective labeling of MCs. On the other hand, the same analysis failed to reveal clear molecular markers for sub-classes of TCs (Supplementary Fig. 5).

Having identified candidate markers for MCs, we sought to test if Cre-recombinase expression driven at these loci would allow MC-specific labeling. Screening several public depositories, we found that a Cre-driver line for *Lbhd2* already exists under a synonymous gene symbol (A230065H16Rik) on GENSAT, a large depository of BAC-mediated transgenic mouse lines (Gong et al., 2007; Heintz, 2004). Since the two independent *Lbhd2-Cre* transgenic lines (*Ra31-Cre* vs. *Ra13-Cre*) show similar recombination patterns, we chose to analyse the line *Ra13-Cre*. As above, we crossed *Ra13-Cre* mice with *Ai14* reporter mice to analyse the pattern of Cre-mediated recombination in the brain (Fig. 3). At P7, red fluorescence was highly selective, showing dense and restricted expression in the cells of the MC layer of the olfactory bulb (Fig. 3A–C; mean number of fluorescent TCs as a proportion of fluorescent cells in the mitral cell layer = 0.09 ± 0.04 for P7; p = 0.18, t-test for mean = 0, t-statistic = 2; n = 3 mice). Correspondingly, labelled dendrites were observed preferentially in the lower portion of the external plexiform layer (fluorescence signal density = 0.20 ± 0.03 for lower external plexiform layer vs. 0.10 ± 0.01 for upper external plexiform layer; p = 0.03, two-sample t-test for equal means, n = 3 mice each), consistent with MCs having dendrites that ramify in the deeper portion of the external plexiform layer. At this developmental stage, red fluorescence was observed only sparsely in the rest of the brain, except for the lateral septum and the dorsomedial nucleus of the hypothalamus. However, in older mice, the residual recombination becomes widespread and is observed throughout the brain. In the olfactory bulb at this stage, while the labeling is still restricted to the projection neurons, substantial number of tufted cells also become labelled (mean number of fluorescent cells in the upper external plexiform layer as a proportion of fluorescent cells in the mitral cell layer = 1.05 ± 0.08 for P21 and 1.33 ± 0.12 for P42). A Cre-driver line that we generated for the second marker candidate, *Pkib*, was deemed unsuitable for MC-specific labeling due to late-onset expression in MCs, as well as a wide-spread recombination in neurons other than MCs (Supplementary Fig. 6).

**Figure 3:**
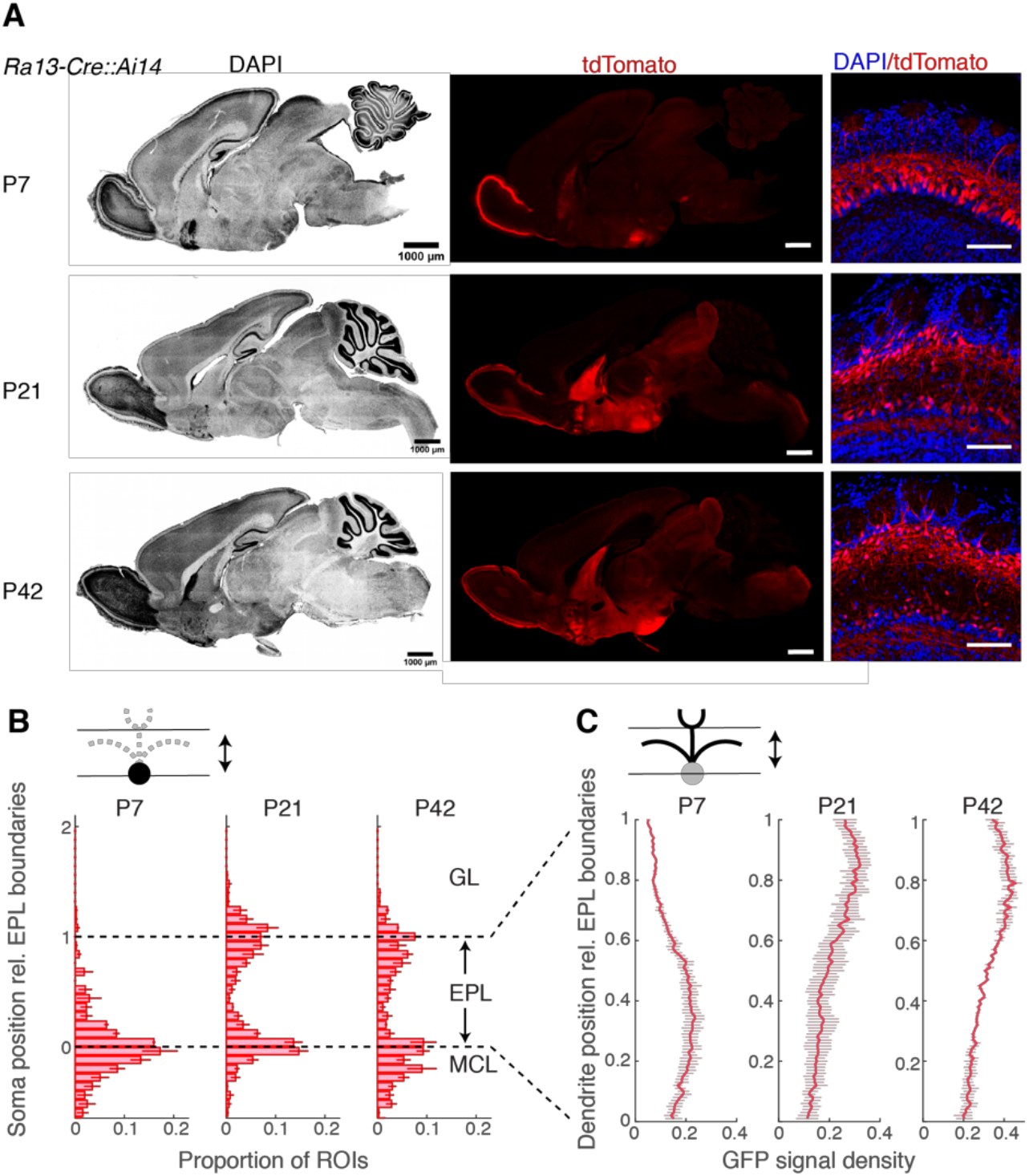
Ra13-Cre∷Ai14 line reveals a developmental accumulation of recombination outside of MCs. (**A**) Sagittal brain sections from example P7 (top row), P21 (middle row) and P42 (bottom row) mice, showing DAPI signal (left column; pseudo-colored in gray scale), corresponding tdTomato (middle column) and DAPI and tdTomato signals for OB (right column). Scale bar = 1 mm for the whole sagittal view, and 100 μm for OB. (**B**) Distribution of somata positions with respect to the external plexiform layer boundaries. N = 3 mice (average of measurements from anterior, ventral and dorsal parts for each animal). (**C**) Distribution of labelled dendrites with respect to the external plexiform layer boundaries.

The developmental accumulation described above makes *Ra13-Cre* unsuitable for investigating MCs in adult mice. However, the *in-situ* hybridization signal for lbhd2 mRNA indicates a clear preference for MCs in the adulthood. Therefore, it is possible that, when the recombination efficiency is calibrated appropriately, a more selective labeling of MCs may be feasible. To this end, we generated a new knock-in line (Fig. 4A) using CRISPR/Cas9, where the variant of Cre-recombinase that is inducible, namely Cre*-ERT2* (Feil et al., 1997), is inserted into the 3′UTR of the lbhd2 gene (the target sequence: 5′-ACCAAGAGGACCTCCAT-3′; Fig. 4A).

**Figure 4:**
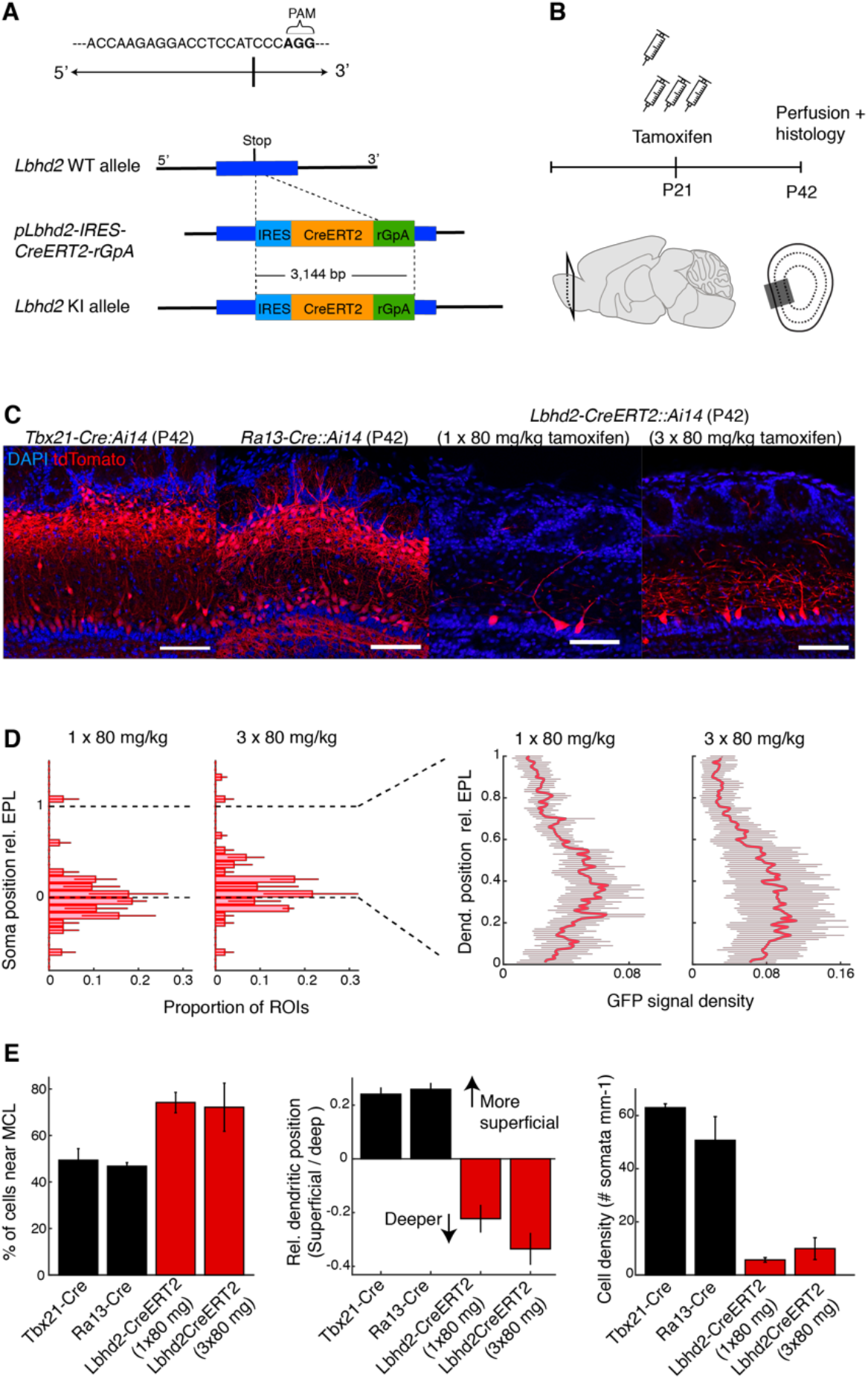
Lbhd2-CreERT2 line achieves MC-specific labeling in the olfactory bulb even in the adulthood. (**A**) Constructs for CRISPR/Cas9-mediated knock-in line. IRES-CreERT2 cassette was targeted to a region immediately following the stop codon of the Lbhd2 gene. (**B**) Schematic of the tamoxifen injection protocol: tamoxifen was injected intraperitoneally at P21, and the recombination pattern in the OB was examined 3 weeks later. Cohorts of mice received injection for either one or three days at 80 mg/kg *per diem*. (**C**) Example recombination patterns. From left: *Tbx21-Cre∷Ai14*, *Ra13-Cre∷Ai14*, *Lbhd2-CreERT2∷Ai14* (one injection) and *Lbhd2-CreERT2∷Ai14* (three injections) mice. Scale bar = 100 μm. (**D**) Summary of labelled structures at P42 for the corresponding mouse lines, showing proportion of cells in the mitral cell layer relative to all labelled cells (left), comparison of labelled dendrites in the superficial vs. deep portions of the external plexiform layer (middle), and density of labelled somata in the mitral cell layer (right). (**E**) Comparison of labeling patterns between *Tbx21-Cre∷Ai14*, *Ra13-Cre∷Ai14* and *Lbhd2-CreERT2∷Ai14* lines. Left: Labelled cells in the mitral cell layer as a percentage of total number of labelled cells. Middle: tdTomato signal density in the upper half of EPL subtracted by the signal density in the lower half of EPL. Right: Number of labelled cells detected per mm of MCL. N = 3 mice per transgenic line for all plots. Mean and s.e.m. shown.

To test if selective labeling is maintained beyond P7 in the new, inducible Cre-driver line, we injected tamoxifen intraperitoneally at P21 in *Lbhd2-CreERT2∷Ai14* mice, and analysed the distribution of red fluorescence to assess the recombination pattern 3 weeks post injection (Fig. 4B, Supplementary Fig. 7). At the lowest dose tested (one injection of 80 mg/kg), the labeling was sparse (average density of labelled MCs = 5.7 ± 0.9 cells per mm), but 74.2 ± 4.4% of labelled somata were located in the mitral cell layer (Fig. 4C,D). The majority of other labelled cells were TCs, save for sporadic labeling in granule cells, which constituted about 1% of the labelled cells. When the dose was increased to three intraperitoneal injections of tamoxifen (at 80 mg/kg *per diem*, over three days), a denser labeling was achieved (mean density = 10 ± 4 cells per mm) while maintaining specificity, indicating that tamoxifen dose can be calibrated to titrate the specificity and density of labeling. Compared to the patterns of recombination observed with existing lines, namely *Tbx21-Cre* and *Ra13-Cre*, overall, the new line achieves a labeling that is substantially more selective for MCs, as measured by the positions of somata (p = 0.016, F = 6.01, 1-way ANOVA; n = 3 mice for *Tbet-Cre* and *Ra13-Cre*, 4 mice for *Lbhd2-IRES-CreERT2*) and dendrites (p < 0.01, F = 59.31, 1-way ANOVA; n = 3 mice for *Tbet-Cre* and *Ra13-Cre*, 4 mice for *Lbhd2-IRES-CreERT2*). To test if AAV-mediated conditional labeling is possible, AAV1-pCAG-Flex-EGFP-WPRE (100 nL) was stereotaxically injected into the dorsal olfactory bulb at a depth of 400 μm below the brain surface in P21 *Lbhd2-CreERT2∷Ai14* mice. Tamoxifen injections overlapped such that the first of the three injections (3 × 80mg/kg, i.p.) occurred immediately after the AAV injection. Three weeks later, labeling pattern was observed, which showed a predominantly MC-selective labeling similar to the pattern obtained with the *Ai14* reporter line (Supplementary Fig. 8).

Having achieved MC-selective labeling with the *Lbhd2-CreERT2* line, we wished to further characterize the properties of the labelled cells in the olfactory bulb, as well as the distribution of labelled fibres in the olfactory cortices (Fig. 5). Specifically, we wished to assess if the labelled MCs are present uniformly in all domains of the olfactory bulb. To this end, confocal images from coronal sections from anterior, middle and caudal levels of the olfactory bulb from *Lbhd2-CreERT2∷Ai14* mice were analysed. This revealed consistent mitral cell layer labeling in all regions of the olfactory bulb, except for the most anterior level, which showed an absence of labeling on the medial side (Fig. 5B–D). In terms of the projection patterns of labelled fibres in the olfactory cortices, we detected red fluorescent fibres as fascicles throughout the antero-caudal extent of the lateral olfactory tract (Fig. 5A,B), as well as thin fibers with bouton-like structures in the molecular layers of olfactory cortices, including in the anterior olfactory nucleus, olfactory tubercle, anterior and posterior piriform cortices (Figure 5E–G). In addition to the anatomical traits, to assess the labelled cells functionally, we loaded a synthetic calcium indicator, Cal-520 dextran by electroporation using a low intensity protocol (Hovis et al., 2010). Using two-photon microscopy in mice anaesthetized with ketamine and xylazine, odour response properties of labelled MCs vs. superficially located TCs were compared. As above, mice were 42 day-old *Lbhd2-CreERT2∷Ai14* ones that had been injected with 3 × 80 mg/kg tamoxifen at P21. Consistent with previous reports (Ackels *et al.*, 2020; Burton and Urban, 2014; Eiting and Wachowiak, 2020; Nagayama et al., 2010), the labelled cells responded to odours more sparsely than TCs (Fig. 6), confirming that the labelled cells likely correspond to MCs. No obvious difference was observed between labeled and unlabeled MCs. Overall, the results here indicate that the new inducible Cre-driver line, *Lbhd2-CreERT2*, achieves a highly specific labeling of MCs in the olfactory bulb.

**Figure 5:**
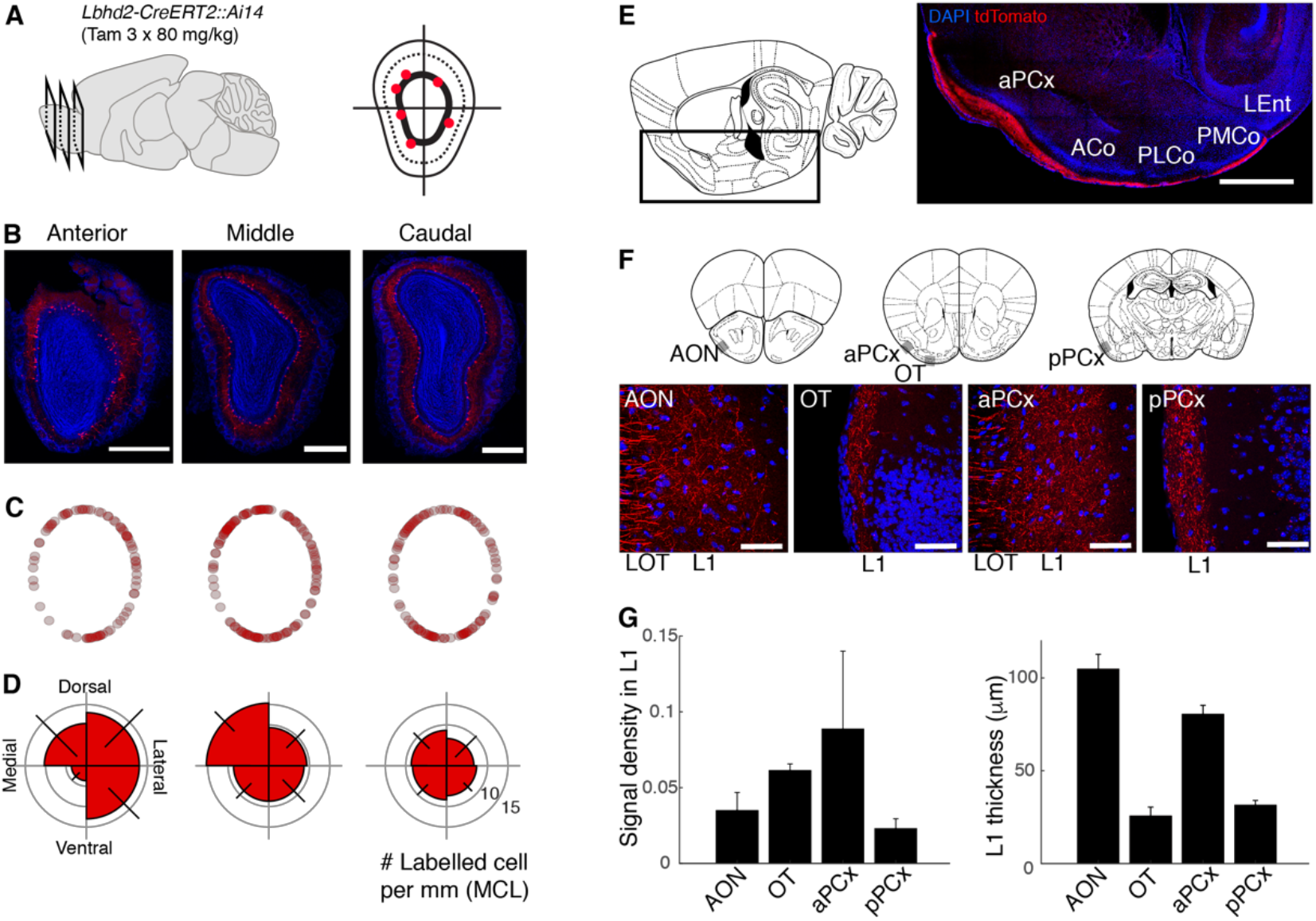
Properties of labelled MCs: OB domain-dependent variations and axon projection patterns. (**A-D**) Distribution of labelled MCs in the OB. (**A**) Schematic of analysis approach: positions of labelled MCs (red dots) were analysed for anterior, middle and caudal levels of the OB of Lbhd2-CreERT2∷Ai14 mice (tamoxifen dose = 3 × 80 mg.kg^−1^). Angular positions were measured relative to the centre of the olfactory bulb. (**B**) Coronal OB images at each antero-posterior level from an example animal. Scale bar = 0.5 mm. (**C**) Positions of labelled MCs from all animals (one dot = one labelled MC), projected on a standardized mitral cell layer position (see Methods). N = 3 mice. (**D**) Polar histograms showing the number of labelled MCs per mm of mitral cell layer for each quadrant. Error bar = s.e.m. (**E-F**) Projection patterns of labelled fibers. (**E**) Image of a sagittal brain section (right) at the medio-lateral plane indicated in the illustration (left), with the imaged location marked by the black outline, showing tdTomato signal present in the lateral olfactory tract (LOT) and the molecular layer for the entire antero-caudal extent of the anterior piriform cortex (aPCx), anterior cortical amygdaloid nucleus (ACo), posterolateral cortical amygdaloid nucleus (PLCo), and posteromedial cortical amygdaloid nucleus (PMCo). Scale bar = 1 mm. (**F**) High magnification (40X objective) of coronal sections taken at the planes shown in illustrations (top). Grey boxes indicate approximate location of images below. Labelled fibres appear as fascicles in the LOT, while densely present thinner, labelled fibers are visible in the superficial, molecular layer (L1) for the anterior olfactory nucleus (AON), olfactory tubercle (OT), anterior and posterior piriform cortices (aPCx and pPCx, respectively). Scale bar = 50 μm. (**G**) Summary quantification of signal density in the molecular layer for the 4 regions (left), and the thickness of the labelled L1 for the corresponding regions (right). N = 3 mice, one image plane each.

**Figure 6:**
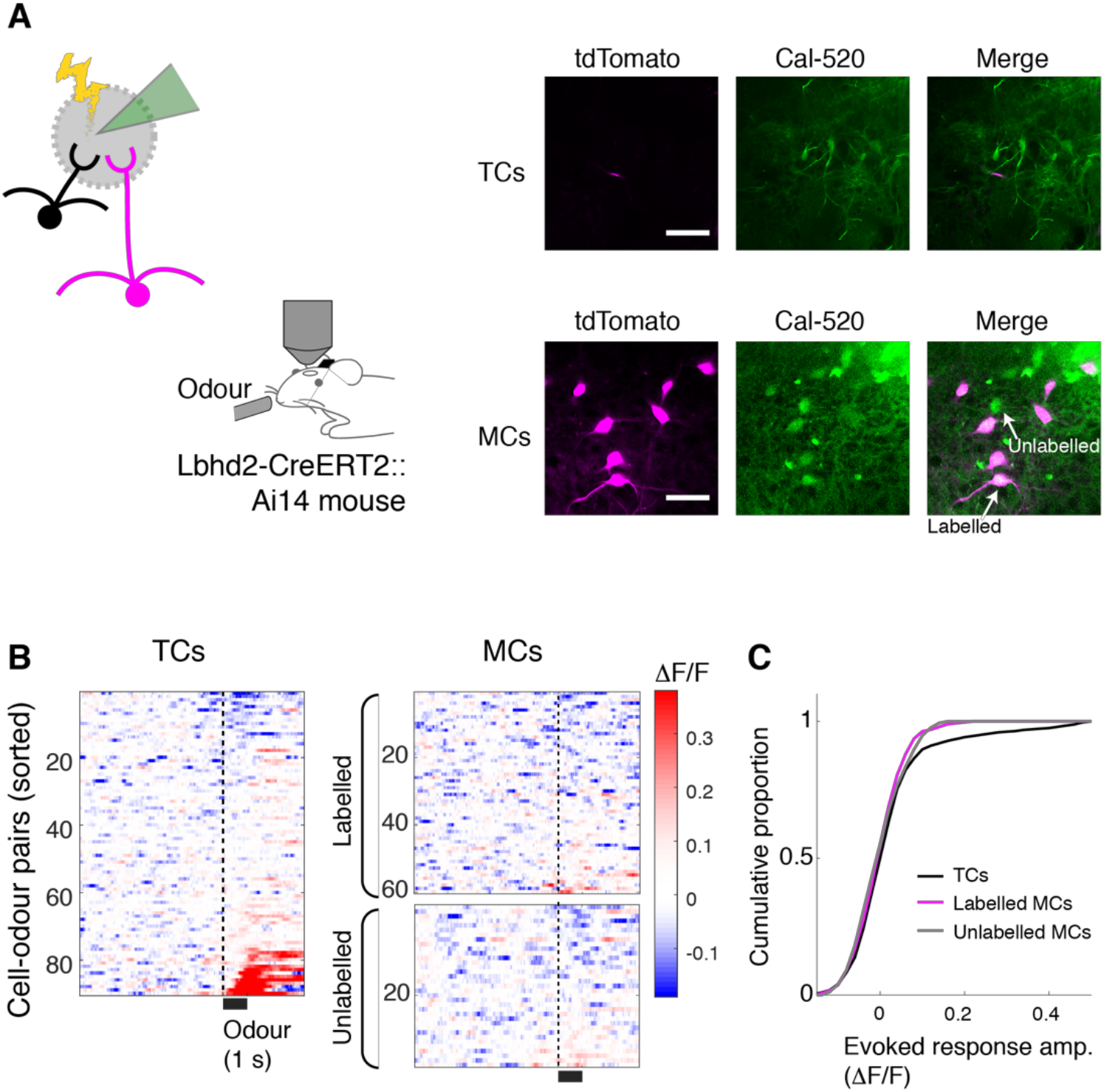
Odour response properties of labelled MCs, in comparison to TCs. (**A**) Left: Schematic showing electroporation of Cal-520 dextran solution in the glomerular layer. MCs were those located ^~^ 300 μm below the brain surface (labeled = red fluorescent cells + green fluorescence, unlabeled = loaded cells without red fluorescence), while TCs were smaller cells located more superficially. Strongly fluorescent cells were excluded from analysis. Scale bar = 50 μm. (**B**) Normalised fluorescence (ΔF/F) from TCs (left) and labeled and unlabeled MCs (right, top and bottom, respectively), shown as colormap (n = 2 mice). Excitatory responses are more prevalent in TCs. Cal-520 has a lower affinity to Ca^2+^ than GCaMP6 variants, which may make hyperpolarizing responses to odours less detectable. (**C**) Cumulative histogram of response amplitude, for TCs (black), labeled MCs (magenta) and unlabeled MCs (grey).

Finally, we examined the recombination pattern in the brain, beyond the olfactory bulb. To assess this, we analysed the distribution of labelled somata in the anterior olfactory nucleus, olfactory tubercle, anterior and posterior piriform cortex, and tenia tacta, as well as other, commonly studied regions, including the thalamus, cerebellum, hippocampus and cerebral cortex. In the *Lbhd2-IRES-CreERT2* mice, for both doses of tamoxifen tested, consistent labeling was observed unexpectedly in a small number of nuclei, including the ventromedial nucleus of the thalamus and the lateral septum, for both doses of tamoxifen (Fig. 7; Supplementary Fig. 9). Compared to the *Ra13-Cre* driver line, the olfactory cortices were devoid of fluorescent cells (Figure 7C; no labelled cells were detected for aPCx, pPCx, AON and OT in *Lbhd2-IRES-CreERT2∷Ai14* mice; in *Ra13*∷*Ai14* mice, mean density of labelled cells = 91.2, 38.5, 490.2, 818.0 labelled cells per mm^2^ for aPCx, pPCx, AON and OT, respectively; p < 0.01, F = 295.04, 2-way ANOVA, with mean densities for *Lbhd2-IRES-CreERT2* groups were significantly different from *Ra13-Cre*; n = 3 mice per group). Thus, the results indicate that the labeling is overall relatively specific to MCs in the whole brain, suggesting that the new inducible Cre-driver line may be suitable for a variety of studies to investigate olfactory processing.

**Figure 7:**
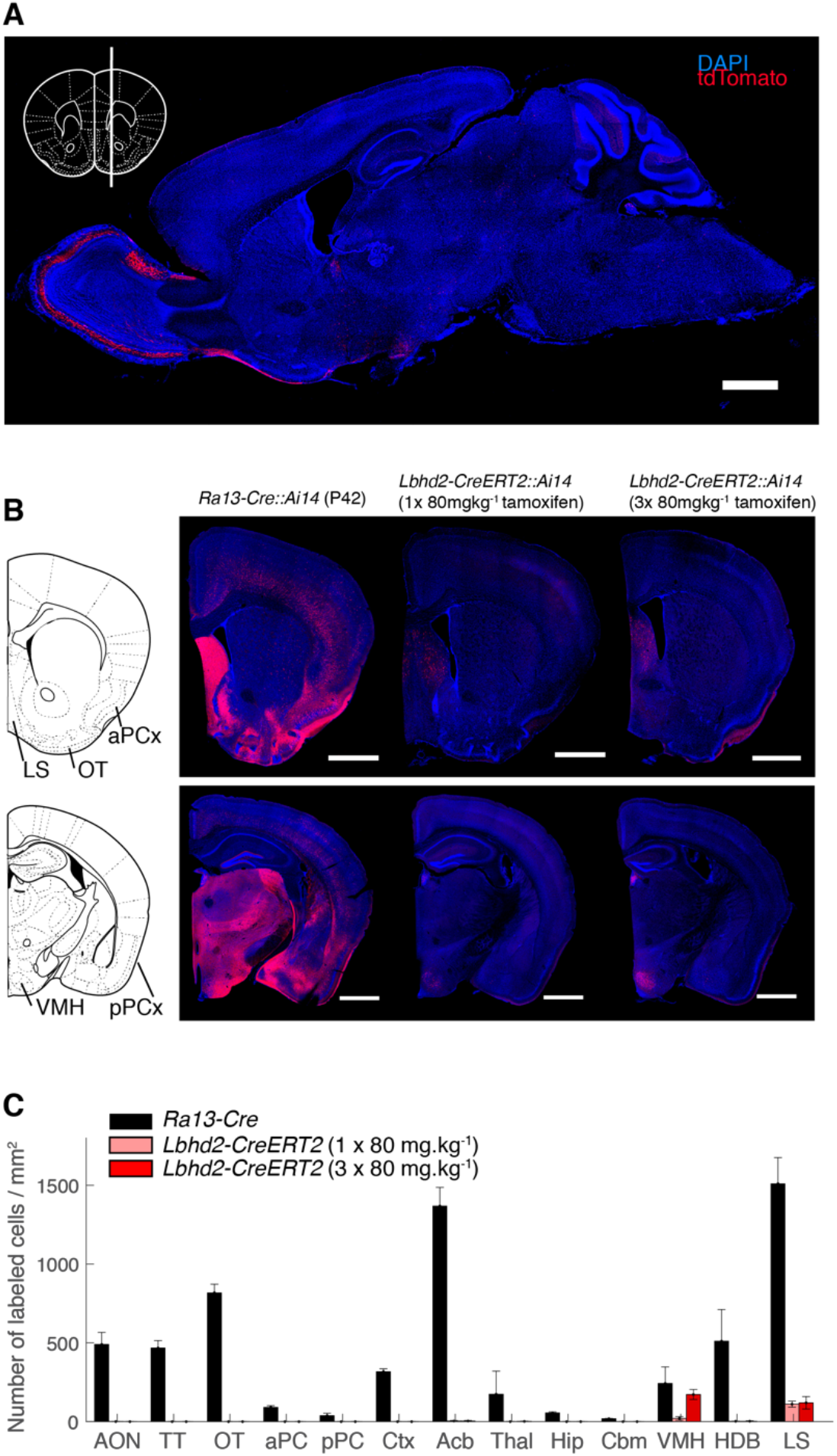
Brain-wide labeling is significantly reduced in Lbhd2-CreERT2 line. (**A**) A sagittal view of an example brain from a P42 Lbhd-CreERT2 mouse, which received 3 doses of tamoxifen (80 mg.kg^−1^) at P21, showing DAPI (blue) and tdTomato (red) signals. Inset shows the medio-lateral plane for the sagittal section. Scale bar = 1 mm. (**B**) Example coronal images from *Ra13∷Ai14* (left), and *Lbhd2-IRES-CreERT2∷Ai14* mice that received 1 × tamoxifen and 3 × tamoxifen doses (middle and right, respectively). Illustrations on the left depict corresponding anatomical borders at this plane. Scale bar = 1 mm. (**C**) Summary showing average density of labelled cells for each anatomical region (n = 3 mice per region). Acb = Accumbens Nucleus (shell); AON = anterior olfactory nucleus; Cbm = cerebellum; Ctx = cerebral cortex; aPCx = anterior piriform cortex; HDB = nucleus of the horizontal limb of the diagonal band; Hip = hippocampus; LS = lateral septum; OT = olfactory tubercle; pPCx = posterior piriform cortex; Thal = thalamus; TT = tenia tecta; VMH = ventromedial nucleus of the hypothalamus.

## Discussion

A wide variety of neuron types that exist in the sensory systems are thought to reflect diverse components for information processing in the brain (Luo *et al.*, 2018; Masland, 2004). Availability of Cre-driver lines have led to a multitude of fundamental insights into unique, cell-type specific contributions to sensory processing and perception (Cruz-Martín *et al.*, 2014; Dhande *et al.*, 2013; Münch et al., 2009; Takahashi et al., 2020). Recent progress in the acquisition, analyses, and applications of large-scale gene expression data have allowed efficient analysis of differences between the cell-types of interest (Birnbaum, 2018; Luo *et al.*, 2018). In this study, we used a publicly available gene expression dataset to discover candidate molecular markers for the key second-order cells of the olfactory system, MCs, which we validated with histology, and finally with new inducible Cre-driver lines generated by CRISPR/Cas9-mediated gene editing. We report that one driver line in particular provides a substantial improvement in the ability to selectively label MCs.

Among the several candidates identified from our differential expression analysis, we found lbhd2 to be the most promising. Specifically, at postnatal day 7, the recombination pattern for the non-inducible, *Lbhd2-Cre∷Ai14* is restricted mainly to MCs in the olfactory bulb. This pattern is consistent with the description on the GENSAT expression database (Gong *et al.*, 2007). Since TCs are already present and located superficially in the external plexiform layer at this stage of development (Mizuguchi et al., 2012), this pattern likely reflects genuine MC specificity in neonatal mice. Despite an increase in the sporadic *Lbhd2*-driven labeling of TCs and other regions of the brain at later stages, the preferential expression in MCs over TCs that persists in the adulthood can be utilized to our advantage. Thus, with our new inducible Cre-driver line, the expression can be targeted selectively to MCs even in the adulthood. It is notable that, despite the fact that many genes were differentially expressed across the two types, markers suitable as genetic tools are harder to identify, especially when selective expression is required across developmental stages. This difficulty may partly be due to the similarity between MCs and TCs, and that cell types are often defined by a combination of genes, rather than a single one (Luo *et al.*, 2018).

In the olfactory bulb, labelled MCs were present in most domains of the olfactory bulb, except for a small patch on the medial, anterior olfactory bulb that showed a curious lack of labeling. Whether or not these correspond to subclasses of MCs, for example those that differ in the glomerular association (Li et al., 2017), or cortical projection patterns (Zeppilli et al., 2020), will be an intriguing future investigation. Outside of the olfactory bulb, labeled somata were sparse if not absent, especially in the areas that MCs target, including in the anterior olfactory nucleus, olfactory tubercle, anterior and posterior piriform. This makes the *Lbhd2-CreERT2* line suitable for investigating the downstream, decoding mechanisms of mitral-specific activity in all these areas, and also when imaging from boutons of MCs (Pashkovski et al., 2020). Beyond these areas, however, we observed a small number of specific areas that showed the presence of labelled somata. The areas include the lateral septum, ventromedial nucleus of the hypothalamus, and the medial amygdala, even at the lowest dose of tamoxifen used. Thus, future studies using this line need to take this into account when interpreting data, in particular for investigating innate, social behavior, which involve these areas (Stowers and Liberles, 2016). However, the fact that only a subset of the nuclei in the pathways are labelled may make this line unexpectedly useful for investigating mechanisms of social behaviours.

This study was aided by publicly accessible data, speeding up discovery. One limitation, if any, in using this dataset for this study may have been the data size, where only a small fraction of olfactory bulb cells expressed *Tbx21*, and even fewer belonged to the putative MC cluster. The relatively small MC cluster size may partly be biological, as glutamatergic neurons comprise heterogenous groups, including those that lack lateral dendrites (Antal et al., 2006; Hayar et al., 2004). MCs comprise only a small proportion of this (Tavakoli et al., 2018). Our histology indicates that superficially located, *Tbx21*-expressing cells are located below the glomerular layer, unlike the *Cck*-expressing population that includes a dense population located more superficially. It should be noted also that a large proportion of glutamatergic and *Cck*-expressing cells were found outside of the *Tbx21*-positive cluster. Some of this latter group may correspond to external tufted cells, which are glutamatergic but lack lateral dendrites (Macrides and Schneider, 1982). It is possible that protocols used to obtain the scRNA-seq data may have been inadvertently biased against large cells with prominent dendrites, such as the filtering step involving a pore size of 30 um (Zeisel *et al.*, 2018).

Despite the need for tamoxifen, this new method for labeling MCs has several advantages over the existing methods. Currently, MC labeling and manipulations are achieved predominantly by depth, birthdate, or retrograde viral expression utilizing the differential projection targets of MCs vs TCs (Economo *et al.*, 2016; Haberly and Price, 1977; Imamura et al., 2011; Rothermel et al., 2013). While this can indeed bias expression patterns, the overlap in somatic and dendritic locations (Schwarz et al., 2018), as well as projection targets (Haberly and Price, 1977; Igarashi *et al.*, 2012) mean that it is not trivial to achieve a perfectly selective labeling. In contrast, our transgenic mouse line described here allows for reproducible and selective labeling of MCs over TCs, with the added advantage that labelled MCs are located throughout the olfactory bulb. Even in imaging applications that can distinguish the cell type based on the soma depth, with the new driver line, it will be possible to investigate the physiology of subcellular compartments, such as the long lateral dendrites, without the need to painstakingly trace back to the somata for cell-type identification. Similarly, investigations of downstream decoding mechanisms, such as one involving precise optogenetic activations of olfactory bulb projections using patterned light stimuli (Chong et al., 2020), may now be done in a cell-type specific manner. Thus, our new tool may bring us closer to understanding how parallel olfactory processing contributes to mechanisms of sensory perception and, ultimately, behaviour.

## Methods

### Gene expression data

Single cell RNA sequence data from the mouse brain was obtained from Zeisel et al (Zeisel *et al.*, 2018) in Loom format. We used the dataset from the level 2 analysis that correspond to the olfactory neurons. The gene expression table represents the expression levels of 27,998 genes in 10,745 olfactory bulb cells. The gene expression level, which counts the number of expressed genes, was transformed into log2(count+1) before analyzing further.

#### Dimensionality reduction

We screened for genes that have higher variability than expected by calculating a log-transformed Fano factor for each gene, as previously described (Li *et al.*, 2017):

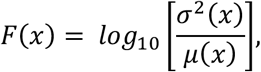

where μ(x) and σ^2^(x) are the mean and the variance of the expression level across cells respectively. Then using the mean expression across different cells, we split the genes into 20 subsets and calculate the Z-score of the Fano factor within each subset:

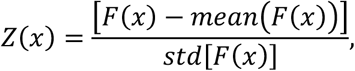

where mean (F(x)) and std[F(x)] are the mean and the standard deviation of F(x) within the subset. Top 500 genes with the highest value of Z(x) were used to cluster the gene expression data. To visualise and cluster the gene expression data corresponding to individual cells in 2D space, we reduce the dimensionality using the principal component analysis (PCA) and t-distributed Stochastic Neighbor Embedding (tSNE; (van der Maaten and Hinton, 2008)). We used top 10 principal components to run tSNE with the following parameters: learning rate = 10, perplexity = 33. The result is shown in Fig. 1. Since many genes have similar expression patterns across different cells, in order to increase the power of PCA and tSNE, we extracted overdispersed genes i.e., the most informative genes.

#### Clustering

Using the 2-dimensional data above, hierarchical clustering algorithm HDBSCAN (Campello *et al.*, 2013) was performed with the following parameters: min_clust_size = 5, min_pts = 13, which indicates which nearest neighbor to use for calculating the core-distances of each point. The cluster with the highest expression level of Tbx21 gene contained 101 cells. Within this group, clustering with HDBSCAN on tSNE space revealed 4 clusters. We then compared the *Cck* expression level across the clusters. We found that 18% (2 of 11) of cells from cluster #4 express *Cck* above the threshold = 3 (log2(count+1), while 64% (58 of 90) from clusters #1, #2 and #3 express *Cck* above the threshold, thus we defined the cluster #4 to be the putative MC cluster.

#### Differential expression analysis and identification of molecular markers

We used the Mann-Whitney U-test to find differently expressed genes. The test works under the assumption that the samples are independent. p values were adjusted using the Benjamini-Hochberg procedure. We screened for significant genes, with the adjusted p value below the threshold = 0.05, where its median expression level in the MC cluster above the threshold = 3 in more than 50% of the cells. Of these, genes that are highly expressed in non-MC clusters were eliminated (cut-off of 10% of cells).

## Animals

All animal experiments have been approved by the animal experiment ethics committee of OIST Graduate University (protocol: 2018-201) and University of Tsukuba. ICR and C56BL/6J mice were purchased from Laboratories International, Inc. (Yokohama, Japan) for generation of transgenic mice, and C56BL/6J from Japan CLEA (Shizuoka, Japan) for subsequent breeding. Ai14 (Madisen et al., 2010) were from The Jackson Laboratory, and Ra13-Cre was from GENSAT project (Gong *et al.*, 2003), via MMRRC (MBP, University of California, Davis).

Generation of *Pkib-IRES-cre and Lbhd2-IRES-CreERT2 mice*: Vector Construction for Knock-in mouse production: The CRISPR target sequence (5′-ATAGCAGCTATGTATTCCTGGGG-3′) was selected for integration of the IRES-Cre sequence just before the stop codon of *Pkib* and *Lbhd2*. The *pX330* plasmid, carrying both gRNA and Cas9 expression units, was a gift from Dr. Feng Zhang (Addgene plasmid 42230) (Cong et al., 2013). The oligo DNAs (Pkib-CRISPR F: 5′-caccATAGCAGCTATGTATTCCTG-3′, and Pkib-CRISPR R : 5′-aaacCAGGAATACATAGCTGCTAT-3′) were annealed and inserted into the entry site of *pX330* as described previously (Mizuno et al., 2014). This plasmid was designated as *pX330- Pkib*. The donor plasmid *pIRES-Cre-Pkib* contained the IRES sequence (Bochkov and Palmenberg, 2006), nuclear translocation signal (NLS)-Cre, and rabbit globin polyadenylation signal sequence. The 1.6-kb 5′-arm (from 1521 bp upstream to 64 bp downstream of *Pkib* stop codon) and the 2.0-kb 3′-arm (from 65 bp downstream to 2,038 bp downstream of *Pkib* stop codon) were cloned into this vector. DNA vectors (*pX330- Pkib* and *pIRES-Cre-Pkib*) were isolated with FastGene Plasmid mini Kit (Nippon genetics, Tokyo, Japan) and filtrated by MILLEX-GV 0.22 μm Filter unit (Merk Millipore, Darmstadt, Germany) for microinjection. Mice were kept in IVC cages under specific pathogen-free conditions in a room maintained at 23.5°C ± 2.5°C and 52.5% ± 12.5% relative humidity under a 14-h light:10-h dark cycle. Mice had free access to commercial chow (MF diet; Oriental Yeast Co., Ltd.., Tokyo, Japan) and filtered water.

### Microinjection and Genomic DNA analyses

The pregnant male serum gonadotropin (5 units) and the human chorionic gonadotropin (5 units) were intraperitoneally injected into female C57BL/6J mice with a 48-h interval and mated with male C57BL/6J mice. We collected zygotes from oviducts in mated females and a mixture of the *pX330- Pkib* (circular, 5 ng/μL) and *pIRES-Cre-Pkib* (circular, 10 ng/μL) plasmids was microinjected into 148 zygotes. Subsequently, survived 137 injected zygotes were transferred into oviducts in pseudopregnant ICR females and 21 pups were obtained.

To confirm the knock-in mutation, the genomic DNA were purified from the tail with PI-200 (KURABO INDUSTRIES LTD, Osaka, Japan) according to manufacture’s protocol. Genomic PCR was performed with KOD-Fx (TOYOBO, Osaka, Japan). The primers (Cre Tm68 F2: 5′-TCTGAGCATACCTGGAAAATGCTTCTGT-3′, and Pkib screening 3Rv: 5′-GTACCAGGAGCTCAAGACAACCTTACCC-3′) were used for checking the 5′ side correct knock-in and the primers (Pkib screening 5Fw: 5′-CTATTTCACAGGTCCAGTTGCTGAAACC-3′, and Cre Tm68 R: 5′-ACAGAAGCATTTTCCAGGTATGCTCAGA-3′) were used for checking the 3′ side correct knock-in. We found 5 of 21 founders carried the designed knock-in mutation. In addition, we checked random integration of *pX330- Pkib* and *pIRES-Cre-Pkib* by PCR with ampicillin resistance gene detecting primer (Amp detection-F: 5′-TTGCCGGGAAGCTAGAGTAA-3′, and Amp detection-R: 5′-TTTGCCTTCCTGTTTTTGCT-3′) and no founder carried the random integration allele.

## Histology

### In situ hybridization

ISH was performed using the RNAscope ISH system (ACDBio, Newark, CA, USA)(Wang et al., 2012).

### Brain extraction

Whole brains were extracted and immediately placed in 4% PFA, dissolved in phosphate buffer (225.7 mM NaH_2_PO_4_, 774.0 mM Na_2_HPO_4_. pH 7.4) at 4°C for 24 hours. Subsequently, the tissues were sunk in DEPC-treated 30% sucrose solution (^~^2 days), then embedded in OCT in a cryomold (Peel-A-Way, Sigma-Aldrich) to be frozen in an ethanol/dry ice bath and stored at −80°C until use.

### Probe design

ISH was carried out using RNAscope (ACDBio, Newark, CA, USA) and probes were produced by ACDBio to be compatible for the procedure. Sequence regions for the *Pkib* and *Ldhd2* probes were selected using the NCBI genetic database. For both probes, regions that were common to all splice variants of each gene were selected. The Pkib probe targeted the region 141-973 bp of the transcript XM_006512605.3. The *Ldhd2* probe targeted the region 138-715 bp of the transcript XM_006516048.1. The Tbx21 probe, which targets the region 893-2093 bp of the transcript NM_019507.2, was already commercially available from ACDBio.

### Hybridisation

On the day of ISH, coronal olfactory bulb sections (20 μm) were cut on a cryostat (Leica CM3050S, Leica Biosystems) at −20°C, washed in RNAse-free PBS (Corning, Manassas, VA, USA), and immediately mounted on glass slides (Superfrost plus). Slides were dried for 30 min at 60°C and post-fixed for 15 min in 4% PFA at 4°C. Slides then underwent *in situ* hybridisation using RNAscope reagents, according to manufacturer’s protocols. Unless otherwise stated, all reagents were provided in the RNAscope kit (RNAScope Intro Pack 2.5 HD Reagent Kit Brown-Mm, Cat no. 322371). Briefly, slides were dehydrated through an ethanol series (75%, 90%, 100%, 100%, Sigma-Aldrich) and endogenous peroxidase activity blocked using provided hydrogen peroxide for 10 min at room temperature. Sections then underwent antigen retrieval by submersion into boiling (^~^98-102°C) 1X Target Retrieval Solution for 5 min and were rinsed in distilled water by submerging 5 times. Subsequently, slides were submerged into 100% ethanol 5 times and air dried. A barrier using an ImmEdge hydrophobic barrier pen was drawn around the sections and left overnight at room temperature to dry. On the following day, slides were treated with Protease Plus and incubated in an oven (HybEZ II System, ACDBio) for 30 min, followed by a series of incubations in the same oven with provided solutions (AMP1-6) to amplify probes (AMP1&3: 30 min 40°C; AMP2&4: 15 min 40°C; AMP5: 30 min RT; AMP6: 15 min RT). After amplification, a DAB reaction was carried out (1:1 mixture of DAB-A and DAB-B solutions) for 10 min at room temperature. Slides were washed by submersion 5 times in 2 changes of distilled water.

### Counterstaining

olfactory bulb sections were immersed in Mayer’s haematoxylin solution (MHS16, Sigma-Aldrich) for 10 mins. Excess stain was washed in distilled water and sections were differentiated by quick submersion in 0.2% ammonium hydroxide in distilled water, followed by washing for 5 minutes in distilled water. Slides were then dehydrated through a series of ethanol for 5 mins each, followed by two 5 min immersions in xylene. Slides were then covered with DPX mountant (06522, Sigma-Aldrich) for histology and left at room temperature to dry before imaging.

## Virus injection

8-10 week old, C57Bl/6J mice were anaesthetised with isoflurane (IsoFlo, Zoetis Japan, Shibuya, Tokyo, Japan) and placed on a stereotaxic frame (Kopf, Tujunga, CA, USA). Carprofen (Rimadyl, Zoetis; s.c., 5mg/kg in saline) was injected subcutaneously for analgesia and aseptic conditions. Fur was trimmed, skin was disinfected with 10% iodine solution before incision. Craniotomy was made bilaterally over centre of the dorsal olfactory bulb (co-ordinates relative to Bregma: AP: 4.8 mm; ML: ± 0.8 mm). 100 nL of AAV1-pCAG-Flex-EGFP-WPRE (Addgene, Watertown, MA, USA) was injected from a pulled glass capillary tube (tip diameter approximately 10 um) at a depth 0.3 mm relative to the brain surface, at a rate of 2 nL every 5 s, using a Nanoject III injector (Drummond Scientific, Broomall, PA, USA). Following injection, the glass capillary was left in place for 1 minute and then slowly withdrawn. The surgical site was then sutured, and mice allowed to recover in a warmed chamber until fully awake, before being returned to their home cage.

## Tamoxifen administration

Tamoxifen solution was dissolved at a concentration of 8 mg/mL in a solvent consisting of 10% ethanol and 90% corn oil (23-0320, Sigma-Aldrich), for once daily injections of 80 mg/kg (10 mL/kg injection volume). Tamoxifen powder (T5648, Sigma-Aldrich) was initially suspended in 100% ethanol and mixed using a vortex mixer to allow partial dissolution. Corn oil was subsequently added to make up solution to the final volume and the solution was heated up to 60°C with agitation on an orbital mixer in an oven, with periodic mixing on the vortex mixer. When fully dissolved (^~^30 min), the final solution was stored at 4°C and used within 1 week.

On the day of injections, the tamoxifen solution was heated up to 60°C with agitation on an orbital mixer for 5 min, and then subsequently mixed on the vortex shaker. After cooling to room temperature, mice were injected intraperitoneally and housed separately from untreated littermates. Mouse weights were monitored carefully throughout the injection period as well as 3 days after the final injection to ensure recovery.

## Two-photon functional imaging

Cal-520 dextran (M.W. ^~^11,000, AAT Bioquest, Sunnyvale USA) was dissolved to 50 mg/mL in Ringer solution comprising (in mM): NaCl (135), KCl (5.4), HEPES (5), MgCl_2_ (1), CaCl_2_ (1.8). Cal-520 dextran solution was electroporated in the OB of P42 *Lbhd2-CreERT2∷Ai14* mice (tamoxifen dose = 3 × 80 mg/kg starting at P21), at a depth ^~^100 μm below the brain surface, under isoflurane anaesthesia. Parameters of electroporation was according to the low intensity protocol described in (Hovis *et al.*, 2010). Subsequently, mice were anaesthetized with ketamine/xylazine and two-photon imaging of dye-loaded TCs and MCs were obtained. MCs were those located ^~^ 300 um below the brain surface (labeled = red fluorescent cells + green fluorescence, unlabeled = loaded cells without red fluorescence), while TCs were smaller cells located more superficially. Strongly fluorescent cells were excluded from analysis.

## Confocal Imaging

Confocal images were acquired on a Zeiss LSM780 confocal microscope with a 10X objective (Zeiss, NA 0.45 Plan-Apochromat) for the whole brain sagittal sections, and 20X objective (Zeiss, NA 0.8 Plan-Apochromat) for the olfactory bulb. Using ZEN 2.3 software (Zeiss), images were taken at a resolution of 1024 × 1024 pixels for a field of view of 850.19 um × 850.19 um (10X) or 425.1 um × 425.1 um (20X objective). To enable comparison and quantification of viral injections, imaging conditions (resolution, gain, laser power, averages) were kept consistent. Sequential laser excitation was used to prevent fluorophore bleed-through. Images were taken throughout the whole rostro-caudal extent of viral spread using the 20× objective. For axonal projection analysis, images were acquired using a Leica SP8 confocal microscope using a 10X (Leica, NA 0.40 Plan-Apochromat) and a 40X (Leica, NA 1.3 Plan-Apochromat) objective. Images were taken at a resolution of 1024 × 1024 pixels per field of view (10X: 1163.64 × 1163.64 um; 40X: 290.91 × 290.91 um) at sequential excitation to prevent fluorophore bleed-trough.

## Image analysis

### In situ hybridisation signal

Images of DAB- and hematoxylin-stained olfactory bulb sections were obtained using a wide-field microscope with a 10X objective in RGB, so that the hematoxylin signal could be separated into the blue channel. Same acquisition setting was used for all sections (Tbx21, Lbhd2 and Pkib signals). Dorsal, ventral, medial and lateral portions of the olfactory bulb at 3 anterior-posterior locations were imaged so that all layers (nerve layer, glomerular layer, external plexiform layer, mitral cell layer and GCL) were captured. To extract the positions of the external plexiform layer boundaries, in ImageJ, a binary mask from the hematoxylin signal (blue) was obtained by setting a threshold and summed along the axis parallel to the olfactory bulb layers (Supplementary Fig. 8). Hybridisation signal (DAB; red channel) was converted into the binary mask, also by setting a single threshold across all conditions. Pixel co-ordinates were normalised such that the boundaries of external plexiform layer were set from 0 – 1, with mitral cell layer being 0. The density of the hybridisation signal was obtained by averaging the binary signal along the axis parallel to the olfactory bulb layers.

### Soma detection and quantification for olfactory bulb

Confocal images (1024 × 1024 pixels corresponding to 425.1 um × 425.1 um) taken with a 20X objective were sampled at anterior, dorsal and ventral locations of the mid-sagittal plane for the tdTomato signal using the *Ai14* reporter line, using DAPI channels to guide sampling, and 10 consecutive planes at a 100 μm interval for the virus injection experiment. Using only the red or green channel for tdTomato labeling and EGFP labelling, respectively, somata were detected manually in ImageJ using ROI manager and their co-ordinates exported into Matlab, without the observer knowing the identity of the mouse. External plexiform layer boundaries were demarcated using only the DAPI signals from images, using a custom-written Matlab routine and the boundary co-ordinates were stored. The soma depths from above were normalised along the external plexiform layer using the boundary co-ordinates, such that mitral cell layer was defined as 0, and the lower boundary of the GL as 1. 1-way ANOVA was used to compare the means, using the *anova1* function in Matlab, and the *multcompare* function with the crucial value tested with Tukey’s honest significant difference criterion for *post-hoc* multiple comparisons. Cells belonging to mitral cell layer were defined as those whose somata are positioned within 30% of the normalised external plexiform layer boundary from the MC layer. This corresponded, on average, to 43.6 μm, which is equivalent to the lengths of two MC somata (Nagayama *et al.*, 2010). Thus, our measure takes into consideration the displaced MCs.

### Dendrite detection and quantification

Images used were the same as those used to detect soma above. To emphasise signals originating from dendrites, which are thin processes, background signal was subtracted from the green or red channel using *Subtract Background* function in ImageJ, with the rolling ball radius set to 5 pixels. Binary masks were created with a single threshold value and the presence of the signal along the normalised external plexiform layer depth at each lateral position was averaged to obtain the density. Dendritic preference index was used to compare the dendritic signal in the upper external plexiform layer vs. lower external plexiform layer, as a proportion of the total dendritic signal detected, calculated as (Signal_density_upper_EPL_ – Signal_density _lower_EPL_)/ (Signal_density_upper_EPL_ + Signal_density _lower_EPL_).

### Analysis of labelled MCs on a standardised coordinate

Labelled MCs from coronal sections (1024 × 1024 pixels, 1.2 um per pixel) were automatically detected in ImageJ by converting the red fluorescence image into binary masks by thresholding and converted into ROIs using the Analyze Particles function (100 – 600 pixels, circularity 0.1-1). Mitral cell layer was delineated using the DAPI channel in Matlab using the drawpolygon function. The line was interpolated, and labelled MCs were projected on the mitral cell layer coordinate. The centre of the olfactory bulb was calculated as the centre of the mitral cell layer coordinates. To pool data across mice, mitral cell layer coordinates were standardised such that it ran from 0 – 2π radians relative to the centre of the olfactory bulb.

### Whole brain somata detection

Positions of somata labelled with tdTomato were automatically detected in the red-channel of the stitched confocal images. To automatically detect the labelled somata, background fluorescence was subtracted using ImageJ’s *Subtract Background* function (100 pixels), then further sharpened to accentuate the somata locally using ImageJ’s *Unsharp* filter with the radius set to 14 pixels, and mask weight set to 0.6. Then a binary mask was obtained by setting a threshold and *Analyze Particles* function was used to detect round objects (size = 70-600 pixels, circularity 0.1 - 1), and detected structures added to the ROI manager, and exported as a list. Using the DAPI signals in the blue channel, boundaries of each nucleus was manually drawn in Matlab using the *drawpolygon* function. Finally, for each anatomical region, all detected soma positions within the boundary was counted using the *inROI* function and normalised by the area to standardise the density of detected cells to be count per mm^2^.

## Acknowledgement

We would like to thank OIST’s Imaging section, sequencing team, and AAALAC-accredited animal facility staff for assistance, and Bernd Kuhn, Sander Lindeman and Gonzalo Otazu for comments on the manuscript. This work was supported by the OIST Graduate University.

## Author contributions

HM and IF conceived the project; AK, CZ, YPH and IF designed the experiments with help from HM and JR; CZ, YPH, TS & JR carried out the experiments; SM, FS and ST generated the transgenic mice; AK and IF analysed the data with help from HM; and IF wrote the manuscript with inputs from all authors.

## Supplementary Figures

**Supplementary Figure 1:**
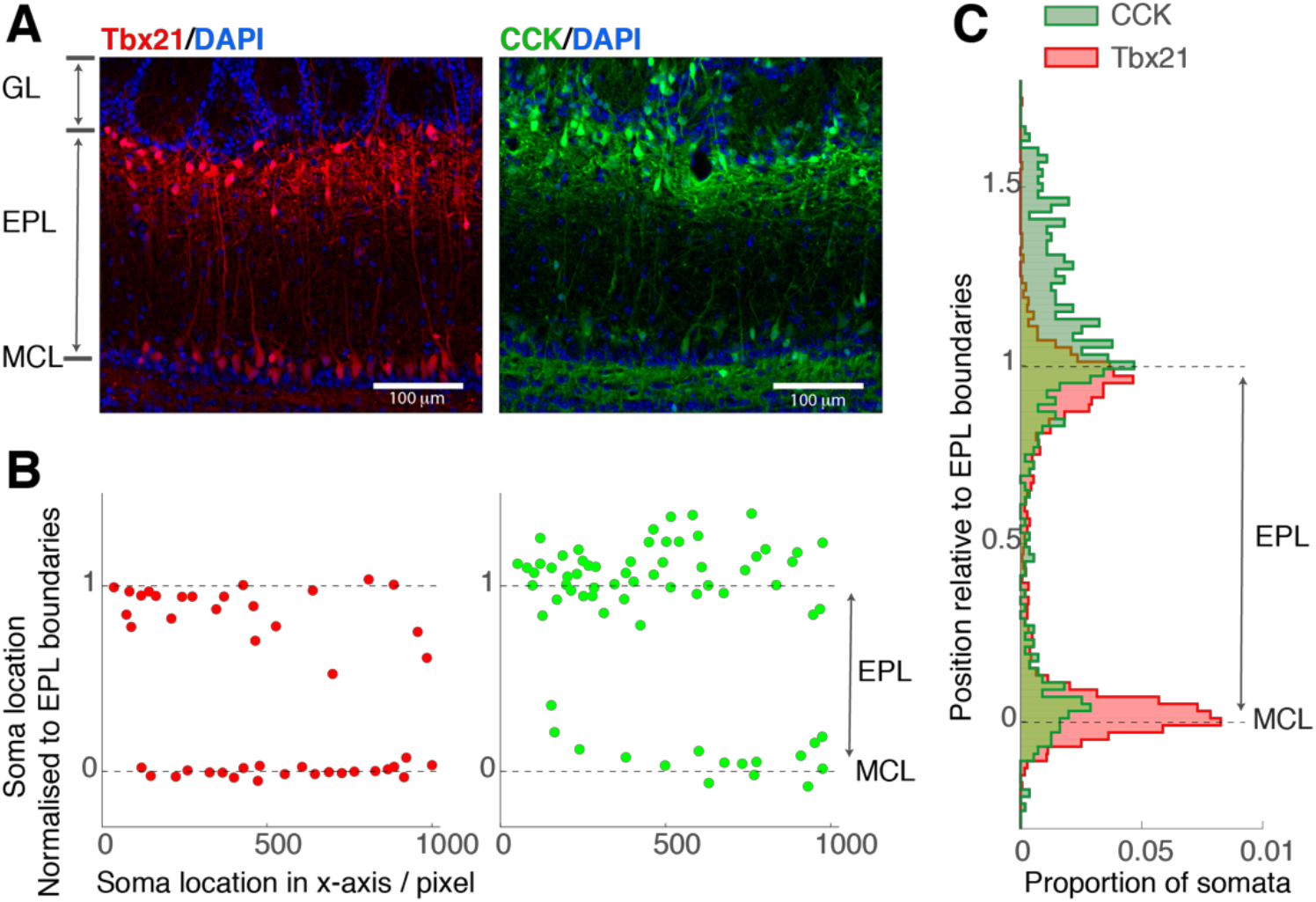
Distribution of Tbet- and CCK-expressing neurons across OB layers. (A) Example confocal images showing OB sections from *Tbet-Cre∷Ai14* (left) and *CCK-IRES-Cre∷Ai14* (right) mice. GL = glomerular layer, EPL = external plexiform layer, MCL = MC layer. Scale bar = 100 um. (B) Soma positions of tdTomato-expressing cells relative to the EPL boundaries, for the images shown in A. EPL depth was normalized so that it run from 0 to 1, with the lower boundary (MCL) corresponding to 0. (C) Summary distribution displayed as the histogram of soma at EPL depths (n = 3 mice).

**Supplementary figure 2:**
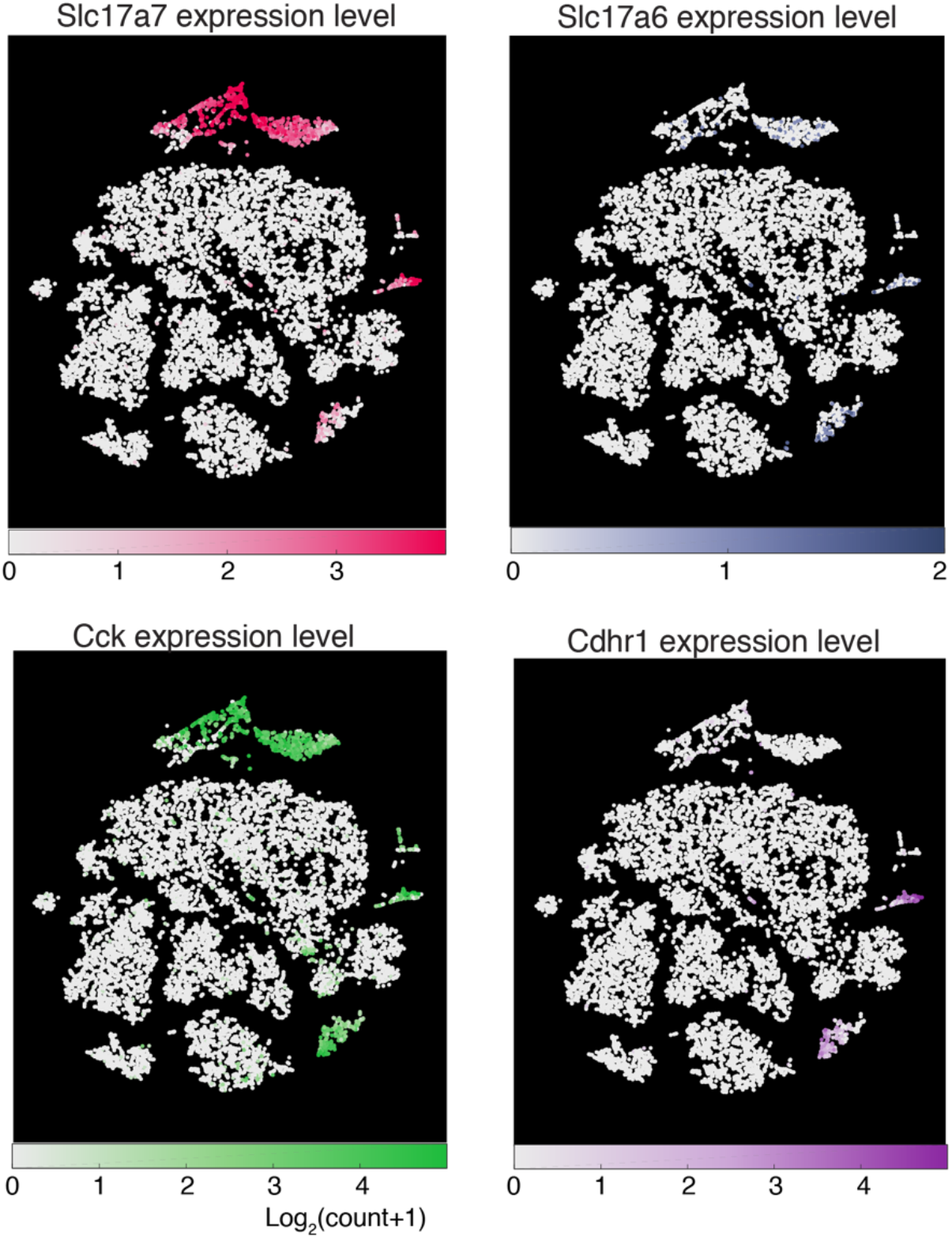
Expression patterns of molecules present in M/TCs. Expression levels of common markers for projection neurons of the OB, namely, VGlut1 (*Slc17a7*), VGlut2 (*Slc17a6*), CCK and *Cdhr1*, for the OB dataset, expressed as log2(count+1) with corresponding colormaps. The t-SNE coordinates are the same as in Figure 1.

**Supplementary figure 3:**
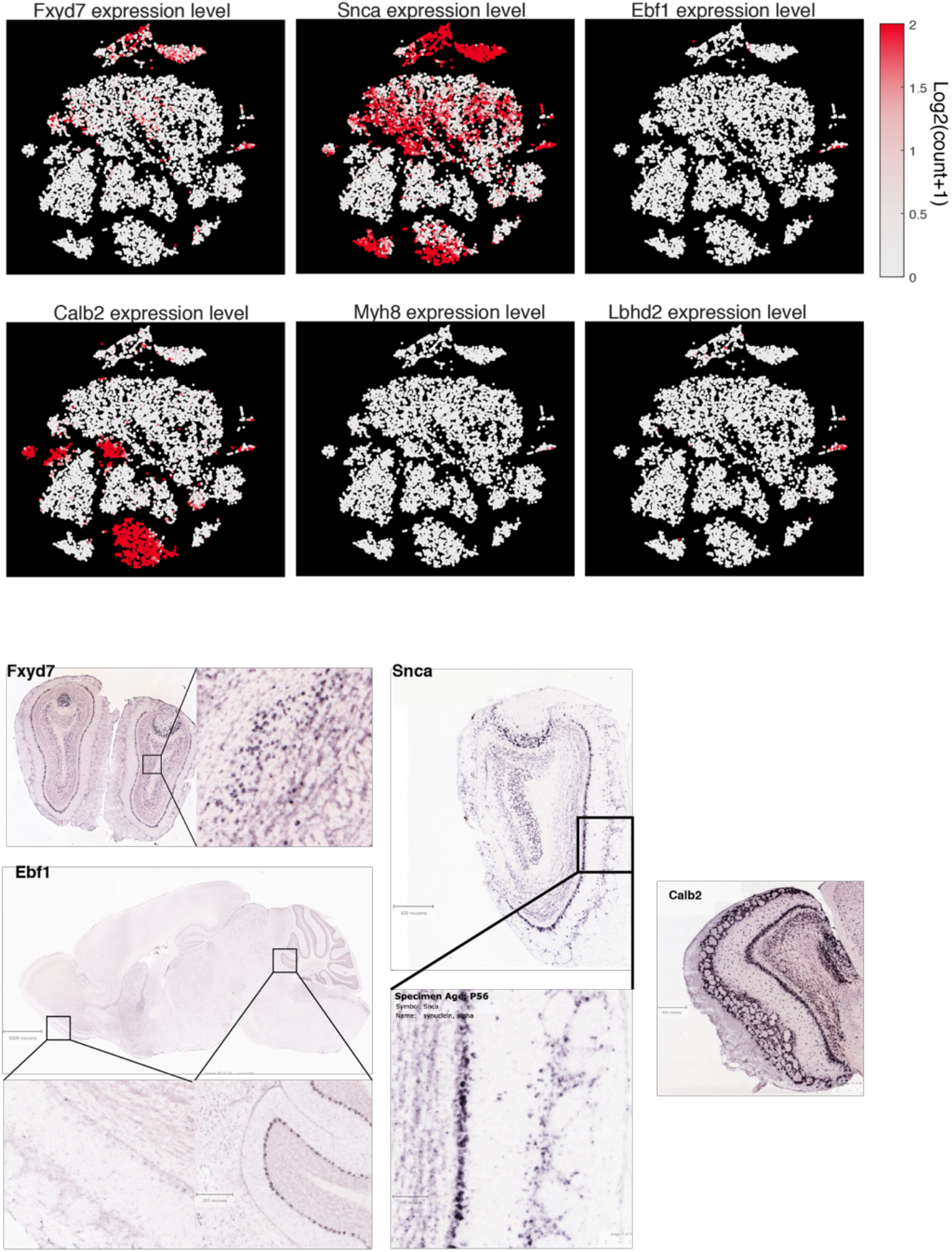
OB-wide tSNE data and *Allen ISH data used to screen some candidate MC markers that were not analysed further.* Differential expression analysis indicates that *Fxyd7*, *Ebf1*, *Snca*, *Calb2*, and *Myh8* are significantly enriched in MCs relative to TCs. To screen candidates, (**A**) the expression pattern (colormap) in the whole OB data were analysed. The same tSNE coordinates as in Fig. 1D are used, with *Lbhd2* expression pattern is shown for comparison. (**B**) Further, ISH database of the Allen Brain Atlas was used to assess the spatial expression patterns. *Fxyd7* seems to be expressed by neurons deep in the granule cell layer as well as superficial cells. *Snca* is expressed by some superficially located neurons as well as some neurons of the anterior olfactory nucleus. *Ebf1* is hardly detectable in the OB even though it is present in the Purkinje cell layer of the cerebellum. Dense *Calb2* hybrdization signal is visible in the glomerular layer, external plexiform layer, mitral cell layer, as well as the granule cell layer. *Myh8* signal was not described in the ISH database but it is a marker for somatostatin positive cells in the subventricular zone (Lim et al., 2018), the source of SST+ve interneurons of olfactory cortices. The expression data in (**A**) shows low levels of Myh8 expression in many neurons outside of the Vglut1- and 2-positive clusters. Image credit: Allen Institute.

**Supplementary figure 4:**
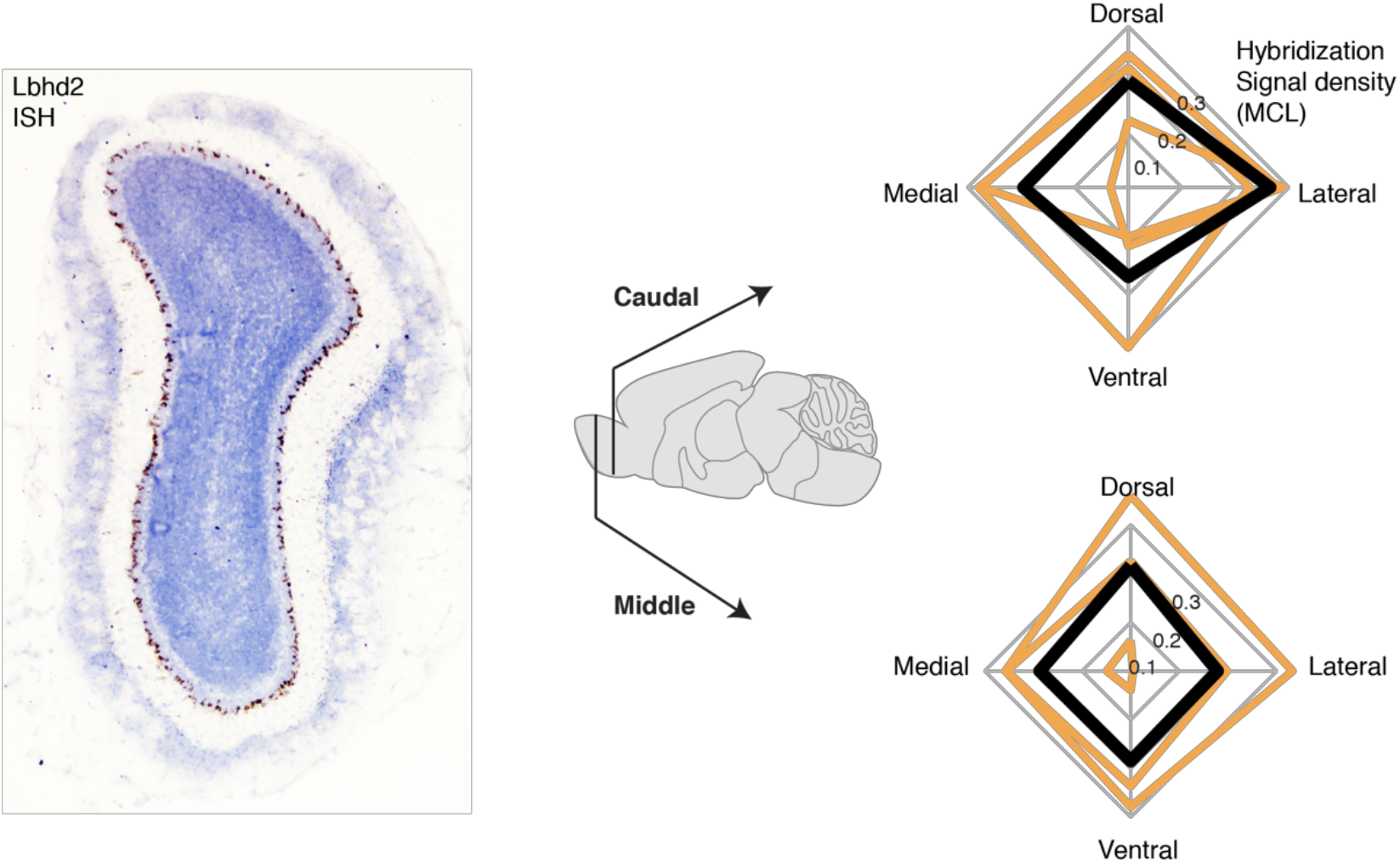
No obvious regional variation in the Lbhd2 expression within the OB. Left: Example coronal section of the OB showing in-situ hybridization signal for Lbhd2, stained with DAB reaction. To quantify the regional variation, average hybridization signal density from the MCL (right) was analysed for dorsal, ventral, medial, and lateral samples taken from middle plane (bottom plot) and caudal plane (top plot) of the antero-posterior axis. Orange lines correspond to data from individual mice, and black lines show the average across the 3 mice.

**Supplementary figure 5:**
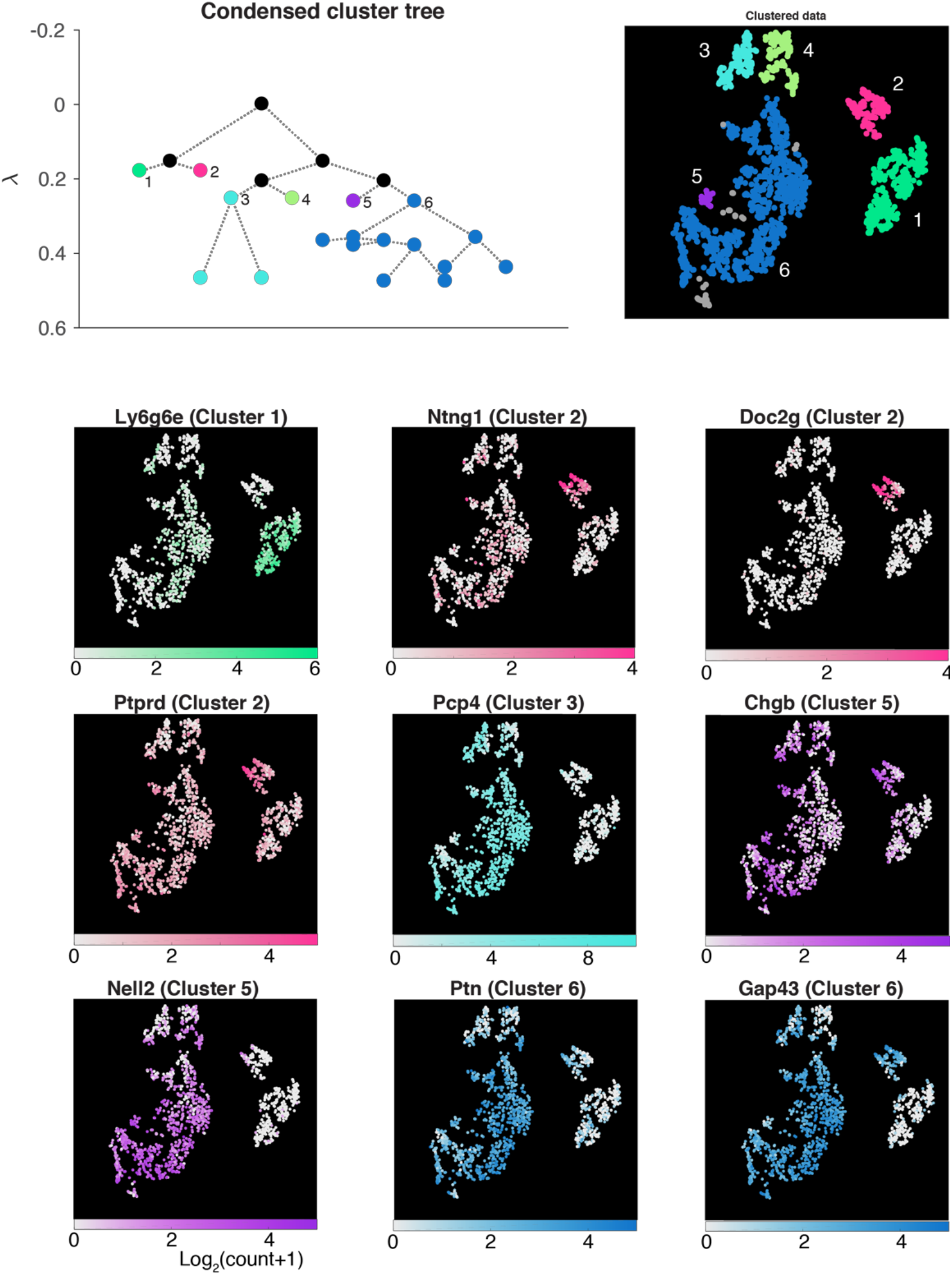
Sub-clustering analysis of CCK-expressing cluster. CCK-expressing population from the OB dataset was further analysed to reveal sub-clusters. For each sub-cluster, candidate marker genes were identified by differential gene expression analysis, where expression patterns from a cluster of interest was compared against all other clusters combined. The expression patterns for each candidate marker, and in which sub-cluster the gene is enriched (in brackets), are shown for all cells in the CCK+ve population, with corresponding colormaps. While Doc2g gene selectively labels the sub-cluster 2 in the TC-dataset, it is a gene that is abundantly expressed by MCs also (Fig. 2).

**Supplementary figure 6:**
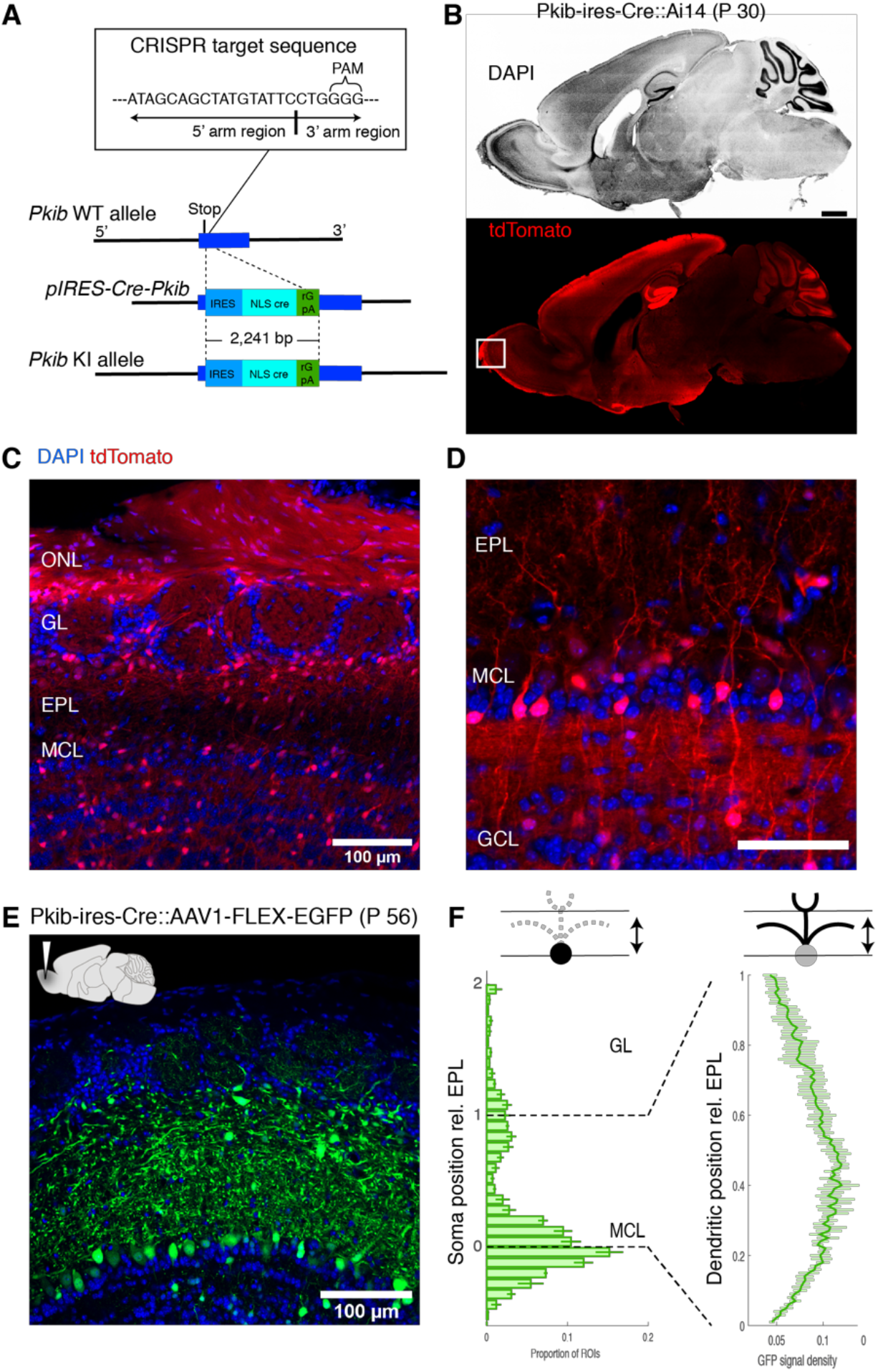
Pkib-ires-cre labels a wide variety of non-MC neurons. (A) Strategy for CRISPR/Cas9 – mediated generation of Pkib-ires-Cre transgenic mouse. CRISPR-target sequence (inset) was just after the stop codon of the Pkib gene. Construct included sequences for IRES, cre-recombinase with a nuclear localization signal, and rGpA. (B) Cre-mediated recombination pattern in a 30-day old Pkib-ires-Cre∷Ai14 mouse. Structures visible in this sagittal plane is revealed by DAPI (top panel), and the corresponding pattern of recombination revealed by tdTomato signal. Scale bar = 0.5 mm. (C) tdTomato expression pattern relative to OB layers from the same animal. ONL = olfactory nerve layer. (D) A higher magnification image. Scale bar = 0.1 mm. (E) A confocal image showing EGFP expression pattern two weeks after an injection of AAV1-flex-EGFP into the MOB in a 42-day old Pkib-ires-cre mouse. (F) Summary of soma location (left) and dendritic signal density (right) relative to the EPL boundaries.

**Supplementary figure 7:**
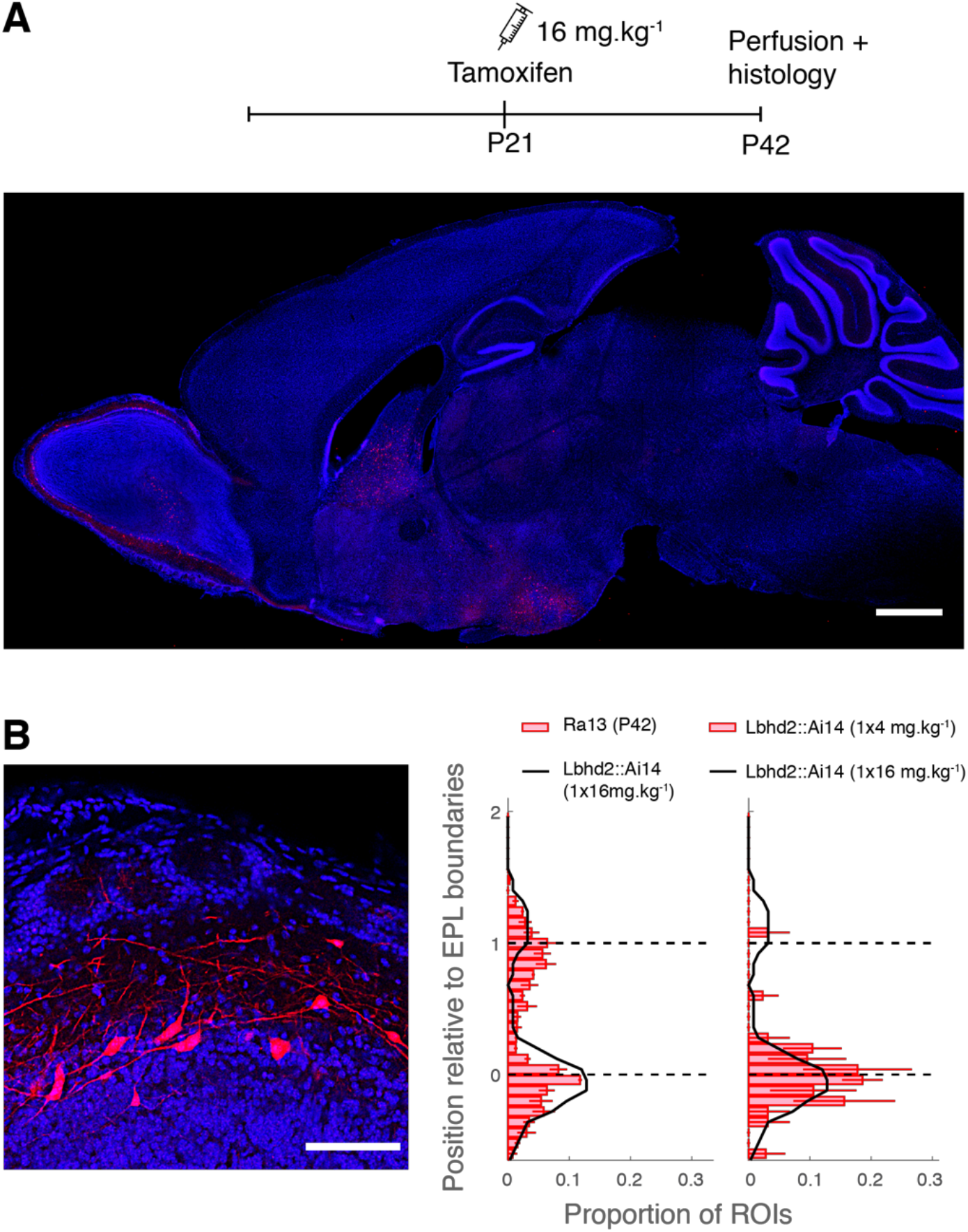
A higher tamoxifen dose increases the density of MCs at the expense of TCs labeling. (**A**) In this example Lbhd2-CreERT2∷Ai14 mouse, tamoxifen (160 mg.kg^−1^) was administered intraperitoneally once, at P21, and tdTomato distribution was determined at P42. Scale bar = 1 mm. (**B**) An example image showing all OB layers, as well as the distribution of soma positions relative to the EPL layers, indicate that while the density of labelled MCs is higher with 1 × 16 mg.kg^−1^ (black line, left and right histograms) than is achievable with a 1 × 80 mg.kg^−1^, but more TCs are labelled. The non-specific labeling in TCs, however, is still less than in Ra13 mice (red bars). Scale bar for the image = 0.1 mm.

**Supplementary figure 8:**
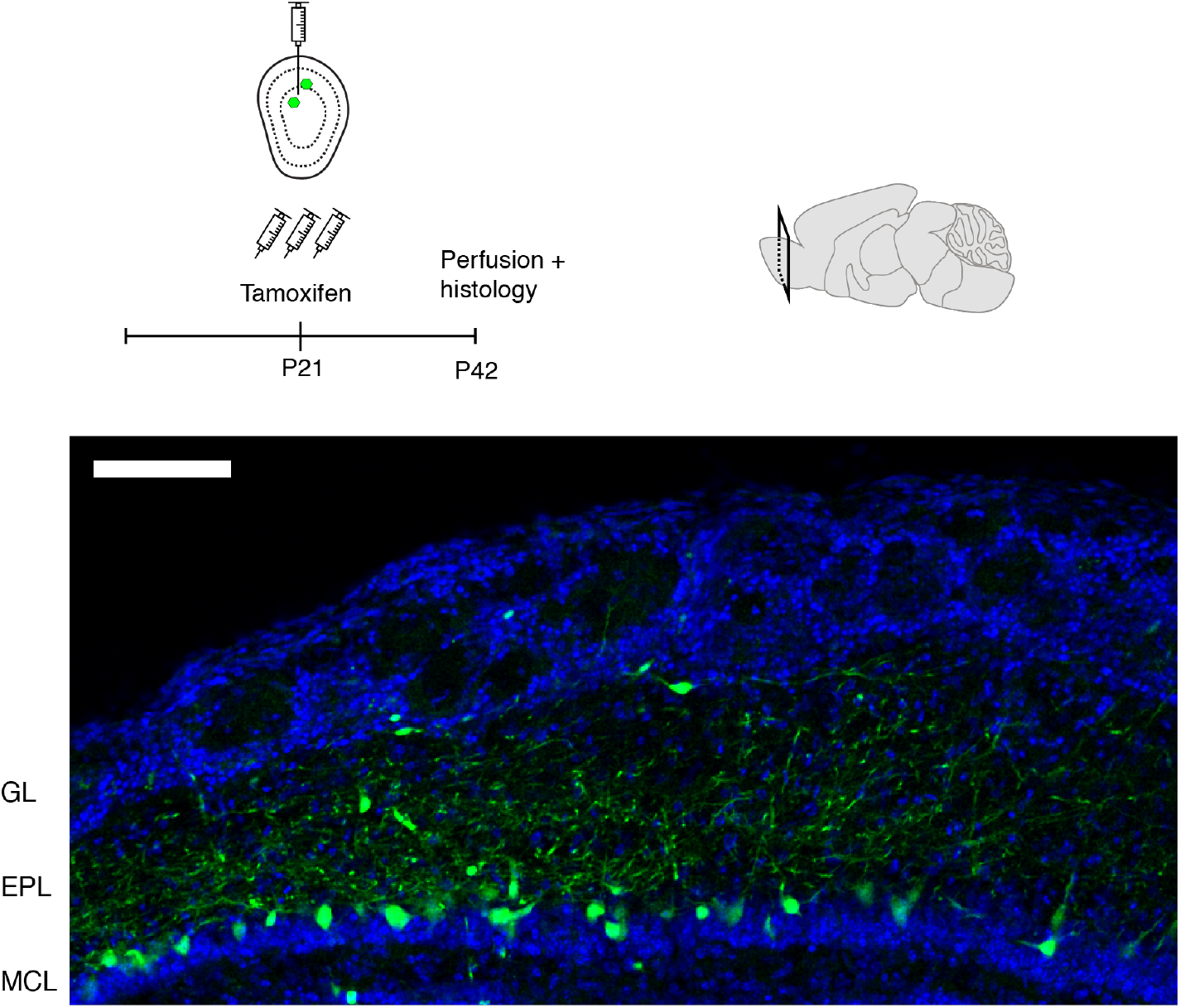
Tamoxifen-induced recombination with AAV-mediated expression. (**A**) In this example Lbhd2-CreERT2∷Ai14 mouse, tamoxifen (3 × 80 mg.kg^−1^) was administered intraperitoneally starting on the day of AAV (AAV1-flex-EGFP) injection in the dorsal OB. Three weeks after the virus injection, at P42, the OB was sectioned and EGFP expression pattern was observed under the confocal microscope.

**Supplementary figure 9:**
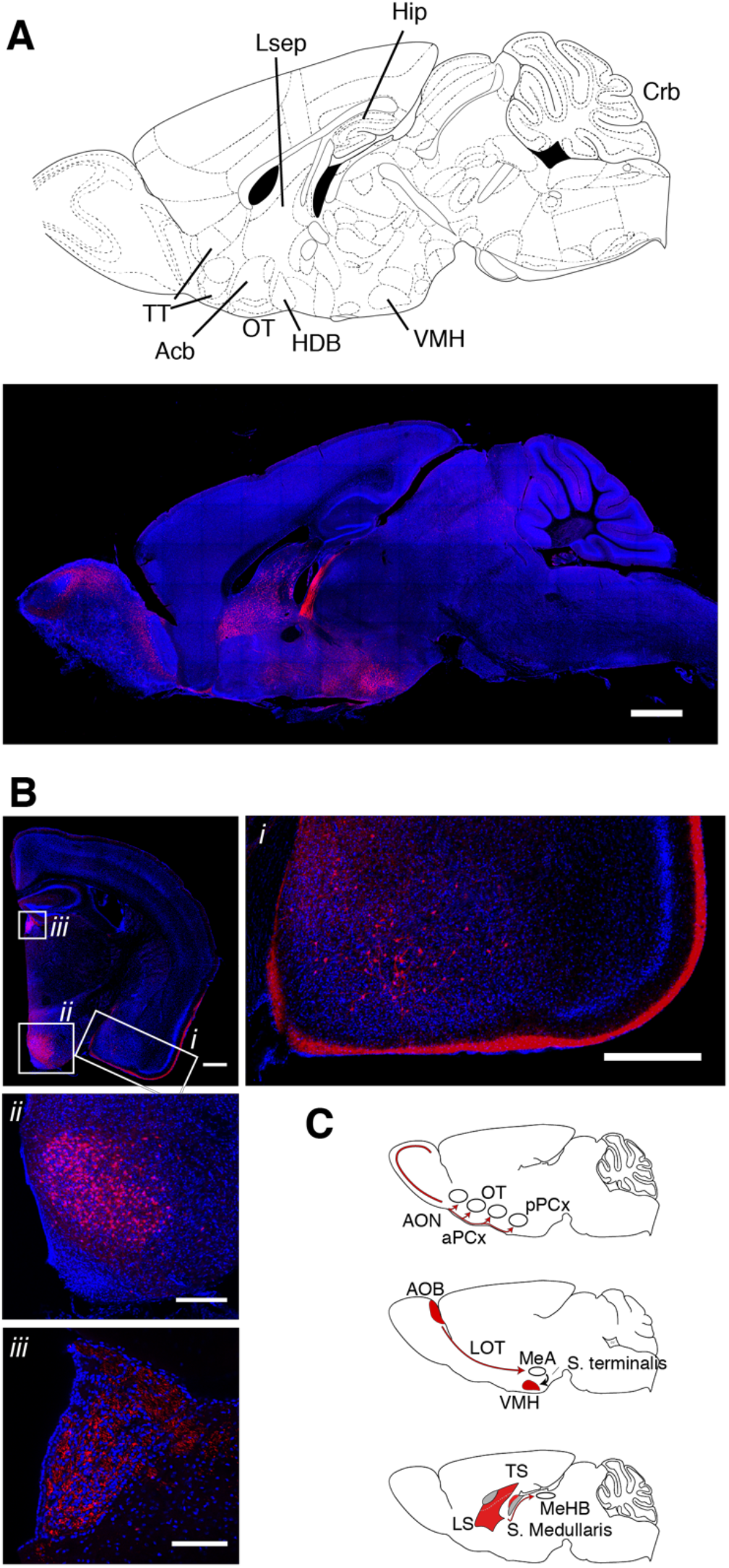
Recombination pattern outside of the OB in Lbhd2-CreERT2 mice. (A) A confocal image at a sagittal plane about 0.36 mm from the midline in a Lbhd2-CreERT2∷Ai14 mouse. Scale bar = 1 mm. (**B**) A coronal view at ^~^1.46 mm posterior to the Bregma, showing (i) labelled cells in the basolateral amygdaloid nucleus, fiber endings in the molecular layer medial amygdaloid nucleus and posterior piriform cortex; (ii) densely labelled somata in the ventromedial nucleus of the hypothalamus and (iii) labelled fibres in the medial habenular nucleus. (C) Summary of labelled structures with respect to distinct pathways; (top) MCs of the main olfactory bulb are labelled, but not their cortical targets. (middle) Principal neurons of the accessory olfactory bulb are labelled. Labelled fibers, but not somata, are visible in the medial amygdaloid nucleus. The target of the medial amygdaloid nucleus, namely, the ventromedial nucleus of the hypothalamus, has densely labelled cells. (bottom) Lateral septum densely contains labelled cells; the output fiber tracts are strongly labelled (stria medullaris), and labelled fibres are clearly visible in the target structure, namely, the medial habenular nucleus.

